# Robust, flexible, and scalable tests for Hardy-Weinberg Equilibrium across diverse ancestries

**DOI:** 10.1101/2020.06.23.167759

**Authors:** Alan M. Kwong, Thomas W. Blackwell, Jonathon LeFaive, Mariza de Andrade, John Barnard, Kathleen C. Barnes, John Blangero, Eric Boerwinkle, Esteban G. Burchard, Brian E. Cade, Daniel I. Chasman, Han Chen, Matthew P. Conomos, L. Adrienne Cupples, Patrick T. Ellinor, Celeste Eng, Yan Gao, Xiuqing Guo, Marguerite Ryan Irvin, Tanika N. Kelly, Wonji Kim, Charles Kooperberg, Steven A. Lubitz, Angel C. Y. Mak, Ani W. Manichaikul, Rasika A. Mathias, May E. Montasser, Courtney G. Montgomery, Solomon Musani, Nicholette D. Palmer, Gina M. Peloso, Dandi Qiao, Alexander P. Reiner, Dan M. Roden, M. Benjamin Shoemaker, Jennifer A. Smith, Nicholas L. Smith, Jessica Lasky Su, Hemant K. Tiwari, Daniel E. Weeks, Scott T. Weiss, NHLBI Trans-Omics for Precision Medicine (TOPMed) Consortium, TOPMed Analysis Working Group, Laura J. Scott, Albert V. Smith, Gonçalo R. Abecasis, Michael Boehnke, Hyun Min Kang

## Abstract

Traditional Hardy-Weinberg equilibrium (HWE) tests (the χ^2^ test and the exact test) have long been used as a metric for evaluating genotype quality, as technical artifacts leading to incorrect genotype calls often can be identified as deviations from HWE. However, in datasets comprised of individuals from diverse ancestries, HWE can be violated even without genotyping error, complicating the use of HWE testing to assess genotype data quality. In this manuscript, we present the Robust Unified Test for HWE (RUTH) to test for HWE while accounting for population structure and genotype uncertainty, and evaluate the impact of population heterogeneity and genotype uncertainty on the standard HWE tests and alternative methods using simulated and real sequence datasets. Our results demonstrate that ignoring population structure or genotype uncertainty in HWE tests can inflate false positive rates by many orders of magnitude. Our evaluations demonstrate different tradeoffs between false positives and statistical power across the methods, with RUTH consistently amongst the best across all evaluations. RUTH is implemented as a practical and scalable software tool to rapidly perform HWE tests across millions of markers and hundreds of thousands of individuals while supporting standard VCF/BCF formats. RUTH is publicly available at https://www.github.com/statgen/ruth.

## INTRODUCTION

Hardy-Weinberg equilibrium (HWE) is a fundamental theorem of population genetics and has been one of the key mathematical principles to understand the characteristics of genetic variation in a population for more than a century (Hardy 1908; Weinberg 1908). HWE describes a remarkably simple relationship between allele frequencies and genotype frequencies which is constant across generations in homogeneous, random-mating populations. Genetic variants in a homogeneous population typically follow HWE except for unusual deviations due to very strong case-control association and enrichment (Nielsen *et al*. 1998), sex linkage, or non-random sampling (Waples 2015).

HWE tests are often used to assess the quality of microsatellite (Van Oosterhout *et al*. 2004), SNP-array (Wigginton *et al*. 2005), and sequence-based (Danecek *et al*. 2011) genotypes. Testing for HWE may reveal technical artifacts in sequence or genotype data, such as high rates of genotyping error and/or missingness, or sequencing/alignment errors (Nielsen *et al*. 2011). It can also identify hemizygotes in structural variants which are incorrectly called as homozygotes (McCarroll *et al*. 2006). Quality control for array-based or sequence-based genotypes typically includes a HWE test to detect and filter out artifactual or poorly genotyped variants (Laurie *et al*. 2010; Nielsen *et al*. 2011).

While HWE tests are commonly and reliably used for variant quality control in samples from homogeneous populations, applying them to more diverse samples remains challenging. When analyzing individuals from a heterogeneous population, the standard HWE tests may falsely flag real, well-genotyped variants, unnecessarily filtering them out for downstream analyses (Hao and Storey 2019). This problem is important since genetic studies increasingly collect genetic data from heterogeneous populations. In principle, HWE tests in these structured populations can be performed on smaller cohorts with homogenous backgrounds (Bycroft *et al*. 2018), and the test statistics combined using Fisher’s or Stouffer’s method (Mosteller and Fisher 1948; Stouffer 1949). However, such a procedure requires much more effort than using a single HWE test across all samples and information that may be imperfect or unavailable.

Here, we describe RUTH (Robust Unified Test for Hardy-Weinberg Equilibrium) which tests for HWE under heterogeneous population structure. Our primary motivation for developing RUTH is to robustly filter out artifactual or poorly genotyped variants using HWE test statistics. RUTH is (1) computationally efficient, (2) robust against various degrees of population structure, and (3) flexible in accepting key representations of sequence-based genotypes including best-guess genotypes and genotype likelihoods. We perform systematic evaluations of RUTH and alternative methods for HWE testing using simulated and real data to explore the advantages and disadvantages of these methods for samples of diverse ancestries.

## MATERIALS AND METHODS

### Unadjusted HWE tests

Consider a study of *n* participants with true (unobserved) genotypes *g*_1_, *g*_2_, ··· *g_n_* at a bi-allelic variant coded as 0 (reference homozygote), 1 (heterozygote), or 2 (alternate homozygote). Represent the best-guess/hard-call (observed) genotypes as 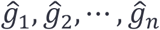. A simple HWE test uses the chi-squared statistic to compare the expected and observed genotype counts assuming no population structure and no genotype uncertainty. The chi-squared HWE test statistic is defined as 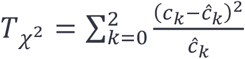 where 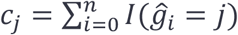 (ignoring missing genotypes), 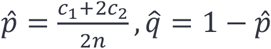, 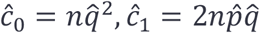, and 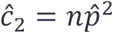. Under HWE, the asymptotic distribution of *T*_*χ*^2^_ is usually assumed to follow 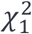 (Rohlfs and Weir 2008). An exact test is known to be more accurate for finite samples, particularly for rare variants (Wigginton *et al*. 2005). HWE tests stratified by case-control status are known to prevent an inflation of Type I errors for disease-associated variants (Li and Li 2008). Widely used software tools such as PLINK (Purcell *et al*. 2007) and VCFTools (Danecek *et al*. 2011) implement an exact HWE test based on best-guess genotypes. We will refer to the exact test as the unadjusted test.

### Existing HWE tests accounting for structured populations

The unadjusted HWE test assumes that the population is homogeneous. If a study is comprised of a set of discrete structured subpopulations, a straightforward extension of the unadjusted test is to (1) stratify each study participant into exactly one of the subpopulations, (2) perform the unadjusted HWE test for each subpopulation separately, and (3) meta-analyze test statistics across subpopulations to obtain a combined p-value using Stouffer’s method (Stouffer *et al*. 1949). More specifically, let *z*_1_, *z*_2_, ···, *z_s_* be the z-scores from HWE test statistics for *s* distinct subpopulations with sample sizes *n*_1_, *n*_2_, ···, *n_s_*. A combined meta-analysis HWE test statistic across the subpopulations is then 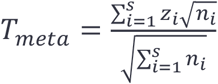, which asymptotically follows a standard normal distribution when each subpopulation follows HWE.

When the population cannot be easily stratified into distinct subpopulations (e.g. intra-continental diversity or an admixed population), a quantitative representation of genetic ancestry, such as principal component (PC) coordinates or fractional mixture over subpopulations, can be more useful for representing each study participant’s genetic diversity (Rosenberg *et al*. 2002; Price *et al*. 2006). HWES takes PCs as additional input to perform HWE tests under population structure with logistic regression (Sha and Zhang 2011), and a similar idea was suggested by Hao and colleagues (2016). However, existing implementations do not support sequence-based genotypes (where genotype uncertainty may remain at low or moderate sequencing depth) or other commonly used formats for genetic array data. A recent method, PCAngsd estimates PCs from uncertain genotypes represented as genotype likelihoods (Meisner and Albrechtsen 2019) and uses these estimates to perform a likelihood ratio test (LRT) for HWE, which is similar to the LRT version of RUTH with differences in computational performance (see below).

### Robust HWE testing with RUTH

Here we describe RUTH (Robust and Unified Test for Hardy-Weinberg equilibrium) to enable HWE testing under structured populations, which is especially useful for large sequencing studies. We developed RUTH to produce HWE test statistics to allow quality control of sequence-based variant callsets from increasingly diverse samples. RUTH models the uncertainty encoded in sequence-based genotypes to robustly distinguish true and artifactual variants in the presence of population structure, and seamlessly scales to millions of individuals and genetic variants.

We assume the observed genotype for individual *i* can be represented as a genotype likelihood (GL) 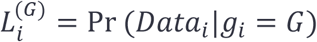, where *Data_i_* represents observed data (e.g. sequence or array), and *g_i_* ∈ {0,1,2} the true (unobserved) genotype. For example, GLs for sequence-based genotypes can be represented as 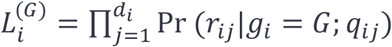 where *d_i_* is the sequencing depth, *r_ij_* is the observed read, and *q_ij_* is the corresponding quality score (Ewing and Green 1998; Jun *et al*. 2012). We model GLs for best-guess genotypes 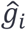 from SNP arrays as 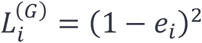, 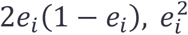 for 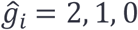 where *e_i_* is assumed per-allele error rate. Imputed genotypes may also be approximately modeled using this framework, but the current implementation requires creating a pseudo-genotype likelihood to describe this uncertainty (see Discussion).

### Accounting for Population Structure with Individual-Specific Allele Frequencies

We account for population structure by modeling individual-specific allele frequencies from quantitative coordinates of genetic ancestry such as PCs, similar to the model (Hao *et al*. 2016). For any given variant, instead of assuming that genotypes follow HWE with a single universal allele frequency across all individuals, we assume that genotypes follow HWE with heterogeneous allele frequencies specific to each individual, modeled as a function of genetic ancestry. Let 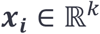 represent the genetic ancestry of individual *i*, where *k* is the number of PCs used. We estimate individual-specific allele frequency *p* as a bounded linear function of genetic ancestry

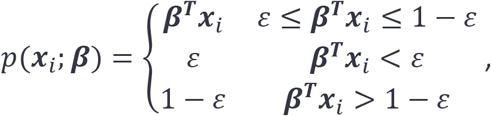

where *ε* is the minimum frequency threshold. We used 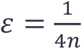 in our evaluation. Even though we used a linear model for *p*(*x_i_*; ***β***) for computational efficiency, it is straightforward to apply a logistic model, which is arguably better (Yang *et al*. 2012; Hao *et al*. 2016).

Let *p_i_* = *p*(***x**_i_*; ***β***) and *q_i_* = 1 – *p_i_* be the individual specific allele frequencies of the non-reference and reference alleles for individual *i*. Under the null hypothesis of HWE, the frequencies of genotypes (0, 1, 2) are 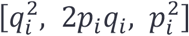. Under the alternative hypothesis, we assume these frequencies are 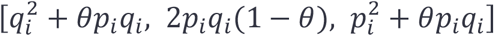 where *θ* is the inbreeding coefficient. This model is a straightforward extension of a fully general model where *p_i_, q_i_* is identical across all samples. Then the log-likelihood across all study participants is

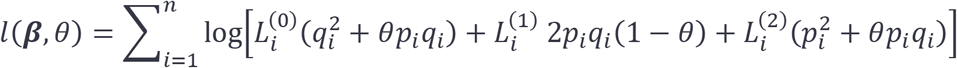

Under both the null (*θ* = 0) and alternative (*θ* ≠ 0) hypotheses, we maximize the loglikelihood using an Expectation-Maximization (E-M) algorithm (Dempster *et al*. 1977). As we empirically observed quick convergence within several iterations in most cases, we used a fixed (n=20) number of iterations in our implementation.

### RUTH Score Test

The score function of the log-likelihood is

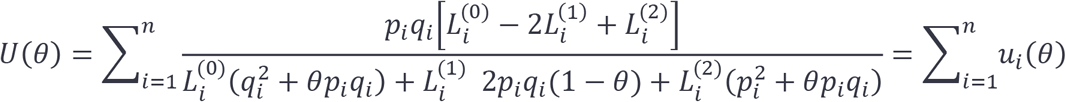

Since 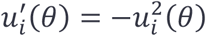, we construct a score test statistic of *H*_0_: *θ* = 0 vs *H*_1_: *θ* ≠ 0 as:

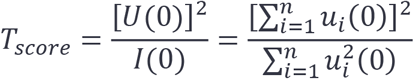

where *l*(0) is the Fisher information under the null hypothesis. Under the null, *T_score_* has an asymptotic chi-squared distribution with one degree of freedom, i.e. 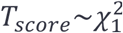. We estimate 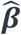 with an E-M algorithm.

### RUTH Likelihood Ratio Test

The log-likelihood function *l*(***β**, θ*) can also be used to calculate a likelihood ratio test statistic:

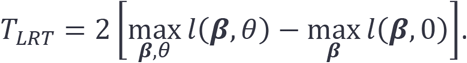

Like the score test, we estimate MLE parameters ***β**, θ* iteratively using an E-M algorithm to test *H*_0_: *θ* = 0 vs *H*_1_: *θ* ≠ 0. Under the null hypothesis, the asymptotic distribution of *T_LRT_* is expected to follow 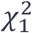. This test is very similar to the likelihood-ratio test proposed by PCAngsd (Meisner and Albrechtsen 2019), except PCAngsd does not re-estimate ***β*** under the alternative hypothesis. In principle, the RUTH LRT should be slightly more powerful due to this difference; we expect the practical difference in power to be small, as deviations from HWE usually do not change the estimates of ***β*** substantially.

### Simulation of genotypes and sequence reads under population structure

We simulated sequence-based genotypes under population structure using the following procedure. First, for each variant, we simulated an ancestral allele frequency and population-specific allele frequencies. Second, we sampled unobserved (true) genotypes based on these allele frequencies. Third, we sampled sequence reads based on the unobserved genotypes. Fourth, we generated genotype likelihoods and best-guess genotypes based on sequence reads.

To simulate ancestral and population-specific allele frequencies, we followed the Balding and Nichols (1995) procedure, except we sampled ancestral allele frequencies from *p* ~ Uniform(0,1) instead of *p* ~ Uniform(0.1,0.9) to include rare variants. For each of *K* ∈ {1, 2, 5,10} populations, we sampled population-specific allele frequencies from 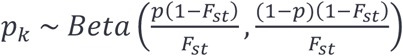, where *k* ∈ {1, “, K}, and *F*_st_ ∈ {.01, .02, .03, .05, .10} was the fixation index to quantify the differentiation between the populations, as suggested by Holsinger (Holsinger 1999) and implemented in previous studies (Holsinger *et al*. 2002; Balding 2003). Because *p_k_* no longer follows the uniform distribution, we used rejection sampling to ensure that 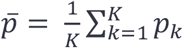 is uniformly distributed across 100 bins across simulations to avoid artifacts caused by systematic differences in allele frequencies.

The unobserved genotype *G_i_* ∈ {0,1,2} for individual *i* ∈ {1, ···, *n_k_*}, belonging to population *k* with sample size *n_k_*, was simulated from genotype frequencies 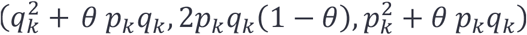, where *q_k_* = 1 – *p_k_* and 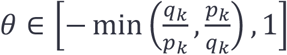 quantifies deviation from HWE; *θ* = 0 represents HWE, while *θ* < 0 and *θ* > 0 represent excess heterozygosity and homozygosity compared to HWE expectation, respectively. In our experiments, we evaluated *θ* ∈ {0, ±.01, ±.05, ±.1, ±.5}. When *θ* was smaller than the minimum possible value for a specific population, we replaced it with the minimum value.

We simulated sequence reads based on unobserved genotypes, sequence depths, and base call error rates. To reflect the variation of sequence depths between individuals, we simulated the mean depth of each sequenced sample to be distributed as *μ_i_* ~ *Uniform*(1, 2*D* – 1), where *D* is the expected depth and *D* = 5 and *D* = 30 representing low-coverage and deep sequencing, respectively. For each sequenced sample and variant site, we sampled the sequence depth from *d_i_ ~ Poisson*(*μ_i_*). Each sequence read carried either of the possible unobserved (true) alleles *r_ij_* ∈ {0,1}, where *j* ∈ {1, ···, *d_i_*}. Given unobserved genotype *G_i_*, we generated 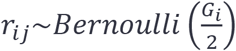, with observed allele *o_ij_* = (1 – *e_ij_*)*r_ij_* + *e_ij_*(1 – *r_ij_*) flipping to the other allele when a sequencing error occurs with probability *e_ij_~Bernoulli*(*ϵ*). We used *ϵ* = 0.01 throughout our simulations (which corresponds to phred-scale base quality of 20) and assumed that all base calling errors switched between reference and alternate alleles.

We then generated genotype likelihoods and best-guess genotypes from the simulated alleles. Let 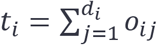 be the observed alternate allele count. The GLs for the three possible genotypes are 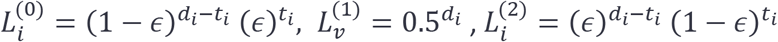. We called best-guess genotypes by using the overall ancestral allele frequency 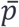 for a given variant as the prior, then calling the genotype corresponding to the highest posterior probability among 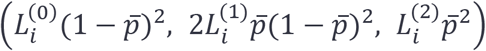 for each sample. For each possible combination of *F_st_*, *K*, and *θ*, we generated 50,000 independent variants across a set of *n* = 5,000 samples with per-ancestry samples sizes 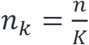.

### Evaluation of Type I Error and Statistical Power

We used different p-value thresholds, *F_st_* values, number of ancestry groups *K*, and average sequencing depth *D* to determine the number of variants significantly deviating from HWE. To evaluate Type I error, we simulated sequence reads under HWE (*θ* = 0) and calculated the proportion of significant variants at each p-value threshold. In RUTH tests, we assumed PCs were accurately estimated using true genotypes unless indicated otherwise. For real data, we summarized ancestral information by projecting PCs estimated from their full genomes onto the reference PC space of the Human Genome Diversity Panel (HGDP) (Li *et al*. 2008) using verifyBamID2 (Zhang *et al*. 2020), similar to the procedure for variant calling in the TOPMed Project, which has already integrated RUTH as part of its quality control pipeline (https://github.com/statgen/topmed_variant_calling).

In all datasets, we evaluated the tradeoff between Type I Error and power for each method using precision-recall curves (PRCs) and receiver-operator characteristic curves (ROCs). In simulated data, we considered variants with θ = 0 to be true negatives and variants with θ = −0.05 to be true positives. In both our 1000G and TOPMed data, we labeled HQ variants as negative and LQ variants as positive.

### Data source

To evaluate our method, we used sequence-based genotype data from the 1000 Genomes Project (1000G) (The 1000 Genomes Project Consortium *et al*. 2015) and the Trans-Omics Precision Medicine (TOPMed) Project (Taliun *et al*. 2019). In both cases, we used a subset of variants from chromosome 20. For 1000G, we started with 1,812,841 variants in 2,504 individuals, with an average depth of 7.0 ×. For TOPMed, we started with 12,983,576 variants in 53,831 individuals, with an average depth of 37.2 ×.

### Application to 1000 Genomes data

To test our method on 1000G data, we first needed to define two sets of variants: one set which is expected to follow HWE, and another set which is expected to deviate from HWE. Unlike simulated data, variants in 1000G are not clearly classified into “true” or “artifactual”, so evaluation of false positives and power is less straightforward. We focused on two subsets of variants in chromosome 20 which serve as proxies for these two variant types. We selected non-monomorphic sites found in both the Illumina Infinium Omni2.5 genotyping array and in HapMap3 (The International HapMap Consortium *et al*. 2010) as “high-quality” (HQ) variants that mostly follow HWE after controlling for ancestry, ending up with 17,740 variants. Similarly, we selected variants that displayed high discordance between duplicates or Mendelian inconsistencies within family members in TOPMed sequencing study as “low quality” (LQ) variants that should be enriched for deviations from HWE even after accounting for ancestry, ending up with 10,966 variants. Among 329,699 LQ variants from TOPMed in chromosome 20, we found that only 10,966 overlap with 1000 Genome samples because likely artifactual variants were stringently filtered prior to haplotype phasing. We suspect that a substantial fraction of these 10,966 LQ variants are true variants since they passed all of the 1000G Project’s quality filters. Nevertheless, we still expect a much larger fraction of these LQ variants to deviate from HWE compared to HQ variants.

We evaluated multiple representations of sequence-based genotypes from 1000G. As 1000G samples were sequenced at relatively low-coverage of 7.0 × on average, best-guess genotypes inferred only from sequence reads (raw GT) tend to have poor accuracy. Therefore, the officially released best-guess genotypes in 1000G were estimated by combining genotype likelihoods (GL), calculated based on sequence reads, with haplotype information from nearby variants through linkage-disequilibrium (LD)-aware genotype refinement using SHAPEIT2 (Delaneau *et al*. 2013). This procedure resulted in more accurate genotypes (LD-aware GT), but it implicitly assumed HWE during refinement. As different representations of sequence genotypes may result in different performance in HWE tests, we evaluated all three different representations - raw GT, LD-aware GT, and GL. In all tests of RUTH using hard genotype calls, we assumed the error rate for GT-based genotypes to be 0.5%, which is representative of a typical non-reference genotype error rate for SNP arrays. We restricted our analyses to biallelic variants. The positions and alleles of 1000G and TOPMed variants were matched using the liftOver software tool (Kuhn *et al*. 2013).

We evaluated all tests as described above. For meta-analysis with Stouffer’s method, we divided the samples into 5 strata, using the five 1000G super population code labels – African (AFR), Admixed American (AMR), East Asian (EAS), European (EUR), and South Asian (SAS). To obtain PC coordinates for 1000G samples, we estimated 4 PCs from the aligned sequence reads (BAM) with verifyBamID2 (Zhang *et al*. 2020), using PCs from 936 samples from the Human Genome Diversity Project (HGDP) panel as reference coordinates. The RUTH score test and LRT used these PCs as inputs, along with genotypes in raw GT, LD-aware GT, and GL formats. For PCAngsd, we used GLs from all variants tested as the input. We limited the analysis to a single chromosome due to the heavy computational requirements of PCAngsd.

### Application to TOPMed Data

We analyzed variants from 53,831 individuals from the TOPMed sequencing study (Taliun *et al*. 2019). These samples came from multiple studies from a diverse spectrum of ancestries, leading to substantial population structure. Using the same criteria as our 1000G analysis, we identified 17,524 high-quality variants and 329,699 low-quality variants across chromosome 20. Since TOPMed genomes were deeply sequenced at 37.2 × (±4.5 ×), LD-aware genotype refinement was not necessary to obtain accurate genotypes. Therefore, we used two genotype representations – raw GT and GL – in our evaluations.

Similar to 1000G, for best-guess genotypes (raw GT), we used PLINK for the unadjusted test. For meta-analysis, we assigned each sample to one of the five 1000G super populations as follows. First, we summarized the genetic ancestries of aligned sequenced genomes with verifyBamID2 by estimating 4 PCs using HGDP as reference. Second, we used Procrustes analysis (Dryden and Mardia 1998; Wang *et al*. 2010) to align the PC coordinates of HGDP panels (to account for different genome builds) so that the PC coordinates were compatible between TOPMed and 1000G samples. Third, for each TOPMed sample, we identified the 10 closest corresponding individuals from 1000G using the first 4 PC coordinates with a weighted voting system (assigning the closest individual a score of 10, next closest a score of 9, and so on until the 10th closest individual is assigned a score of 1, then adding up the scores for each super population) to determine the super population code that had the highest sum of scores, and therefore best described that sample. In this way, we classified 15,580 samples as AFR, 4,836 as AMR, 29,943 as EUR, 2,960 as EAS, and 716 as SAS. Among these samples, 94.5% had the same super population code for all 10 nearest 1000G neighbors. To evaluate the RUTH score test and LRT for both raw GT and GL, we used 4 PCs estimated by verifyBamID2 (Zhang *et al*. 2020), consistent with the method applied for the 1000G data.

### Impact of Ancestry Estimates on Adjusted HWE Tests

We examined the effect of changing the number of PCs used as input for RUTH tests by using 2 PCs as opposed to 4 PCs. We also evaluated the impact of using different approaches to classify ancestry when adjusting for population structure with meta-analysis. By default, our analysis classified the 1000 Genomes subjects into 5 continental super populations based on published information (The 1000 Genomes Project Consortium *et al*. 2015). For TOPMed, the best-matching 1000 Genomes continental ancestry was carefully determined using the PCA-based matching strategy described above. However, in practice, ancestry classification may be performed with a coarser resolution (Jin *et al*. 2019). To mimic such a setting, we used k-means clustering on the first 2 PCs of our samples to divide individuals into 3 distinct groups, and performed meta-analyses based on this coarse classification for both 1000G and TOPMed data.

### Software and data availability

RUTH is available at https://github.com/statgen/ruth. Genotype data from 1000G is available from the International Genome Sample Resource at https://www.internationalgenome.org. TOPMed data is available via a dbGaP application for controlled-access data (see https://www.nhlbiwgs.org for details).

## RESULTS

### Simulation: Effect of Genotype Uncertainty

To evaluate the impact of genotype uncertainty, we first compared tests in the absence of population structure (i.e. single ancestry). For the unadjusted test, we used only best-guess genotypes (GTs). For PCAngsd, we used only genotype likelihoods (GLs). For RUTH score and likelihood ratio tests, we used both.

Using GLs over GTs substantially reduced Type I errors in HWE tests, especially in low-coverage data (Figure 1A-C). For example, the standard HWE test based on GTs resulted in a 229-fold inflation (22.9%) at p < .001 (Figure 1B, Table S1), a threshold which allows the evaluation of Type I error with reasonable precision with 50,000 variants (50 expected false positives under the null). GT-based RUTH-Score and RUTH-LRT tests showed similar inflation. When GLs were used instead of best-guess genotypes, RUTH-Score and RUTH-LRT had Type I errors close to the null expectation (.001 for RUTH-Score and .0012 for RUTH-LRT). PCAngsd, which also accounts for genotype uncertainty (Meisner and Albrechtsen 2019), had similar performance. The severely inflated Type I errors with best-guess genotypes can largely be attributed to high uncertainty and bias towards homozygote reference genotypes in single site calls from low-coverage sequence data, resulting in apparent deviations from HWE. For high-coverage sequence data, inflation of Type I error with GTs was substantially attenuated; inflation nearly disappeared when using GLs (.004 for RUTH-Score and .002 for RUTH-LRT; Figure 1D-F).

**Figure 1.**
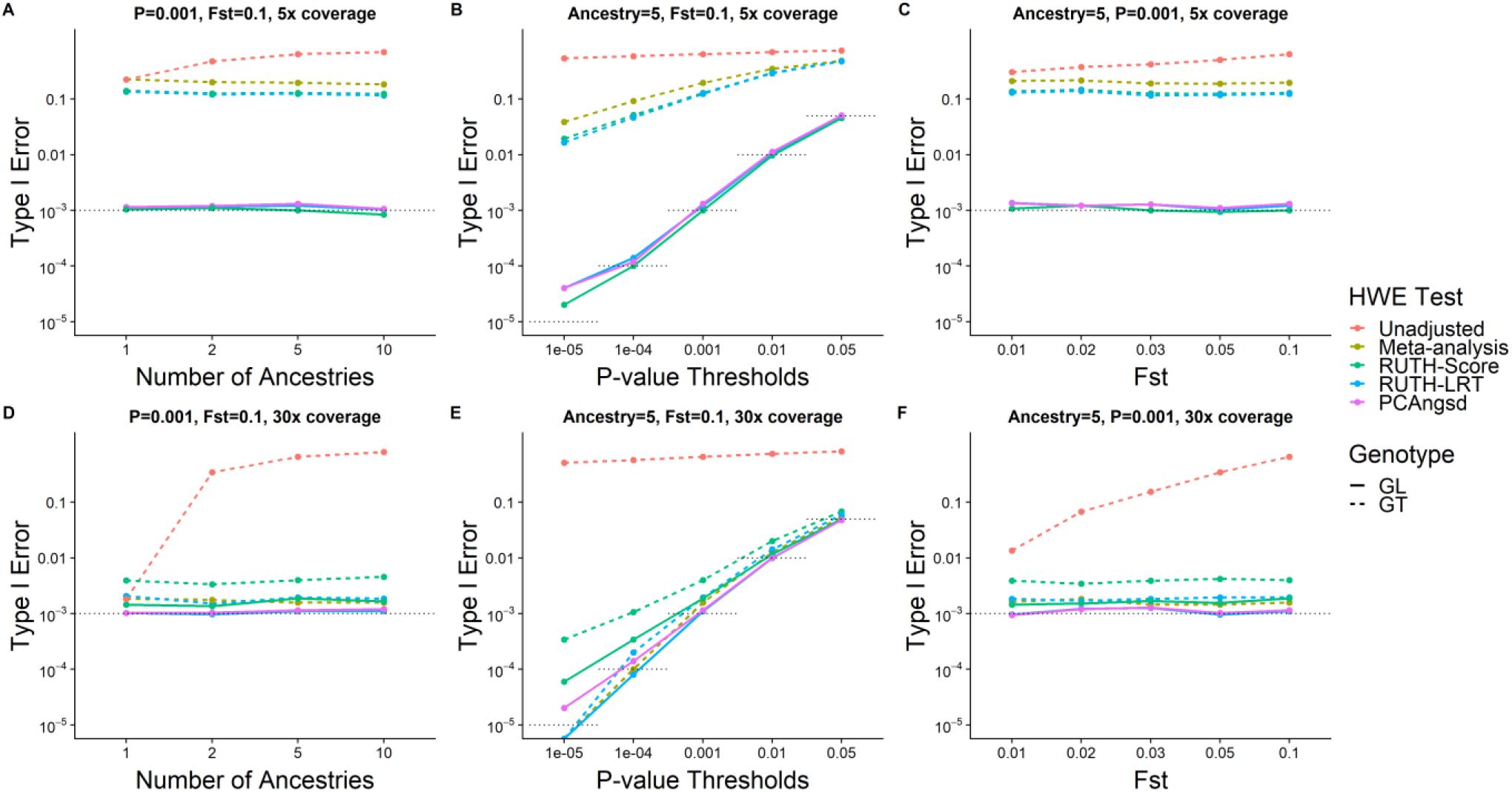
Evaluation of Type I Errors between various HWE tests on simulated genotypes. Under each combination of simulation conditions (number of ancestries, sequencing coverage, and fixation index), we simulated 5,000 samples with 50,000 variants that follow HWE within each of the subpopulations and determined the Type I error performances of different HWE tests based on the proportion of variants labeled as having significant p-values. Five HWE tests – (1) Unadjusted HWE test (Wigginton *et al*. 2005) implemented in PLINK-1.9 (Purcell *et al*. 2007) using hard genotypes, (2) meta-analysis using Stouffer’s method across ancestries using hard genotypes (GT), (3) RUTH test using hard genotypes, (4) RUTH test using phred-scale likelihood (GL) computed from simulated sequence reads, and (5) PCAngsd (Meisner and Albrechtsen 2019) – were tested under HWE with various parameter settings. Gray dotted lines indicate targeted Type I Error rates. Top panels (A-C) represent results from shallow sequencing (5x), and the bottom panels (D-F) represent results from deep sequencing (30x). Using GL-based genotypes resulted in Type I Error rates closer to the targeted rate than using GT-based genotypes across different numbers of ancestries (A, D), P-value thresholds (B, E), and fixation indices (C, F). The difference is especially large for low-coverage genotypes.

Next, we evaluated the power to identify variants truly deviating from HWE at various levels of inbreeding coefficient (θ). For low-coverage sequence data, we skip interpretation of power of GT-based tests owing to their extremely inflated false positive rates. All GL-based tests behaved similarly, achieving ~19-21% power at p < .001 with moderate excess heterozygosity (θ = −0.05) (Figure 2B, Table S1). For high-coverage sequence data, the power of GL-based tests at the same p-value threshold increased to ~56-60%, comparable to corresponding GT-based tests. Interestingly, the unadjusted GT-based test showed much lower power than RUTH and PCAngsd tests under excess heterozygosity (θ < 0) while demonstrating much higher power with excess homozygosity (θ > 0). Upon further investigation, we observed that the tests behave very differently for rare variants for which an asymptotic approximation performs poorly.

**Figure 2.**
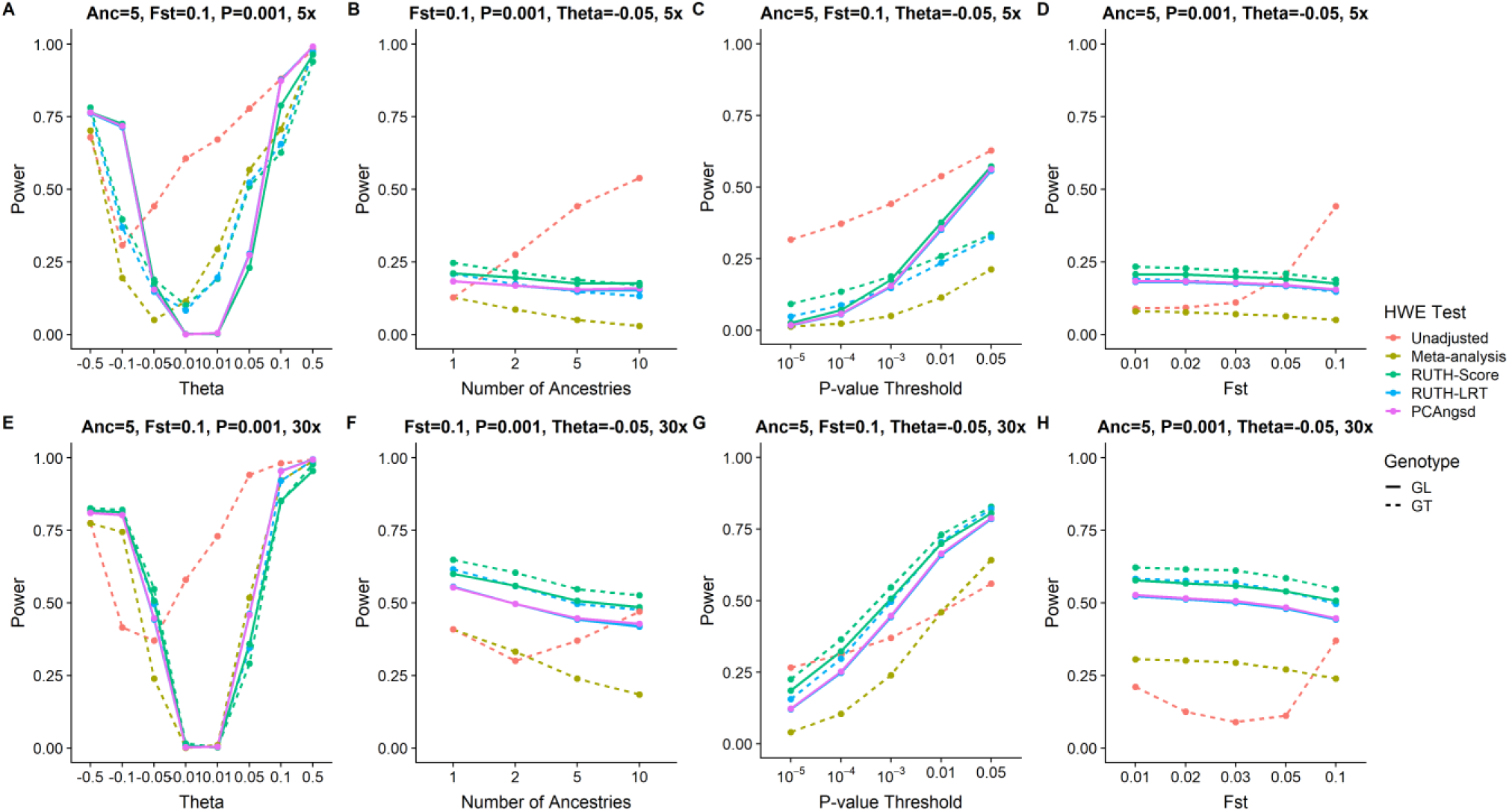
Evaluation of power between different HWE tests on simulated genotypes. Under each combination of simulation conditions (number of ancestries, sequencing coverage, fixation index, and deviation from HWE), we simulated 50,000 variants for 5,000 samples and evaluated the ability of different HWE tests to find the variants significant. Unless otherwise specified, the default simulation parameters are 5 ancestries, with F_ST_=.1, P-value threshold=.001, and Theta=-0.05. Tests that can find a larger proportion of significant variants are considered more powerful. Five HWE tests – (1) Unadjusted HWE test (Wigginton *et al*. 2005) implemented in PLINK-1.9 using hard genotypes (2) RUTH test using hard genotypes, (3) RUTH test using phred-scale likelihood (PL) computed from simulated sequence reads, (4) meta-analysis using Stouffer’s method across ancestries using hard genotypes, and (5) PCAngsd (Meisner and Albrechtsen 2019) - were tested for variants deviating from HWE with various parameter settings, for low coverage (A-D) and high coverage (E-H) data. (A, E) Theta controls the degree of deviation from HWE, with negative values indicating excess heterozygosity and positive values indicating heterozygote depletion. The high Type I Error rates in GT-based tests (Figure 2) lead to those methods appearing to have higher power in some scenarios. The unadjusted test suffers from this problem the most. GL-based methods have slightly lower powers than GT-based methods in exchange for a much better controlled Type I error rate. This pattern mostly holds across different numbers of ancestries (B, F), p-value thresholds (C, G), and fixation indices (D, H). Meta-analysis had the lowest power in the presence of excess heterozygosity.

We also generated precision-recall curves (PRC) and receiver-operator characteristic (ROC) curves to better understand the tradeoff between the Type I errors and power under moderate excess heterozygosity (θ = -.05) (Figure S1C-D). Again, accounting for genotype uncertainty resulted in better empirical power and Type I error, especially for low-coverage data, for which, at an empirical false positive rate of 1%, GL-based tests had 41-45% power, as opposed to 4-10% for GT-based tests. For high-coverage data, GL-based tests had 1-2% greater power than GT-based tests at the same false positive rate. These results suggest that ignoring genotype uncertainty in HWE tests is reasonable for high-coverage sequence data.

### Simulation: Impact of Population Structure on HWE Test Statistics

As expected, the unadjusted HWE test had substantially inflated Type I errors under population structure based on the Balding-Nichols (1995) model (Figure 1, Table S1). Even for an intra-continental level of population differentiation (F_ST_ = .01), the Type I errors at p < .001 were inflated 13.5-fold even for high-coverage data. With an inter-continental level of differentiation (F_ST_ = .1), we observed orders of magnitude more Type I errors across different simulation conditions. This inflation is expected to increase with larger sample sizes, suggesting that adjustment for population structure is important even if a study focuses on a single continental population.

One simple approach to account for population structure is to stratify individuals into distinct subpopulations to apply HWE tests separately (Bycroft *et al*. 2018), and meta-analyze the results (Figure 3B). Type I errors were appropriately controlled with this approach in high-coverage but not low-coverage data, likely due to unmodeled genotype uncertainty (Figure 1, Table S1). Instead of classifying individuals into distinct subpopulations, RUTH incorporates PCs to jointly perform HWE tests (Figure 3C). For both low- or high-coverage data, GL-based RUTH tests and PCAngsd showed well-controlled Type I errors, while GT-based tests showed slight (high-coverage) or severe (low-coverage) inflation.

**Figure 3.**
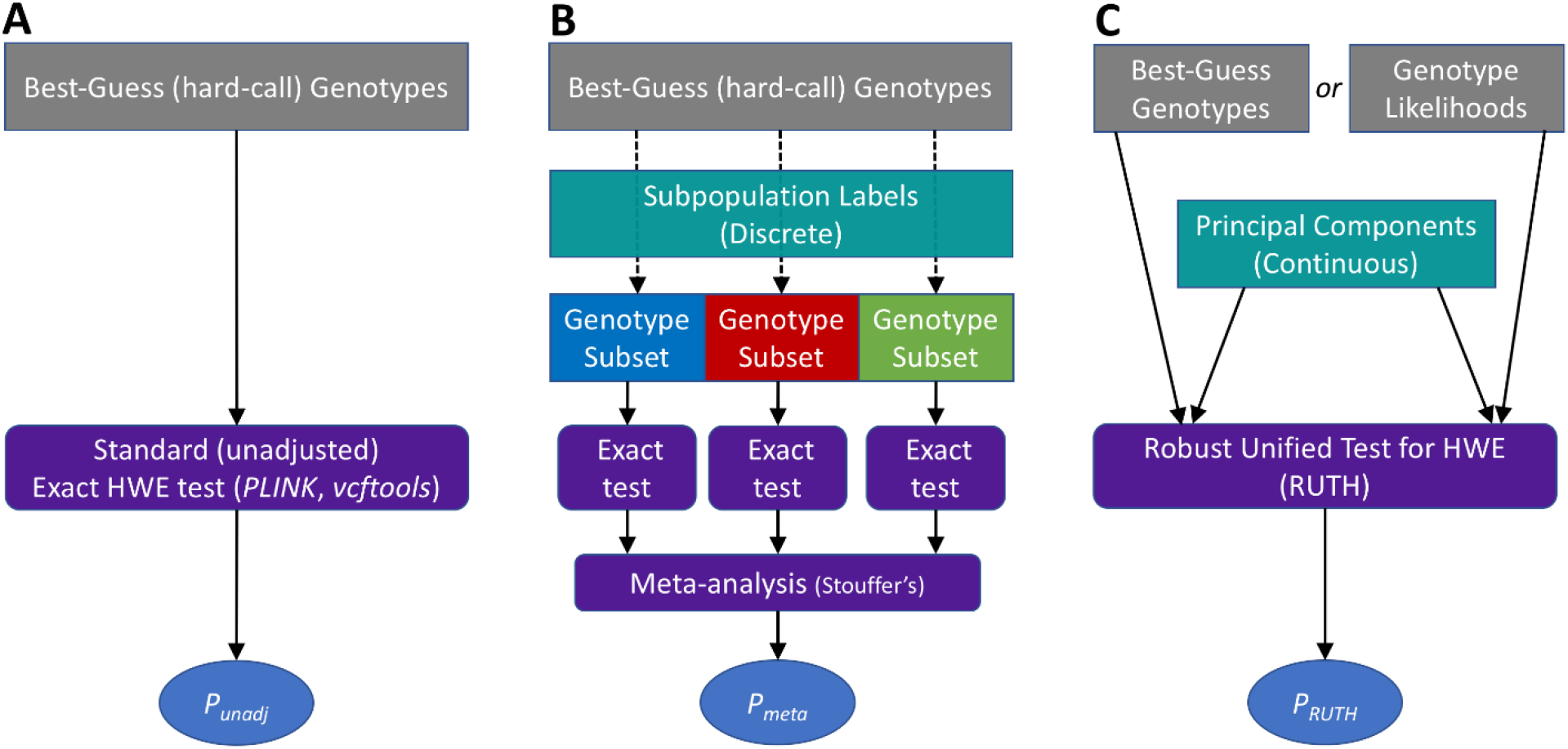
Schematic diagrams of different methods to test HWE under population structure. Three different methods to test HWE under population structure are described. (A) In the standard (unadjusted) HWE test, all samples are tested together using best-guess genotypes. This test does not adjust for sample ancestry. (B) In a meta-analysis of stratified HWE tests, the samples must first be categorized into discrete subpopulations, determined a priori based on their genotypes or self-reported ancestries. Next, standard HWE tests (based on best-guess genotypes) are performed on each of these subpopulations. Then, the resulting HWE statistics are converted into Z-scores and combined in a meta-analysis using Stouffer’s method, with the sample sizes of the subpopulations as weights. (C) In our proposed method (RUTH), either best-guess genotypes or genotype likelihoods can be used as input for HWE test. We assume that the genetic ancestries of each sample are estimated a priori, typically as principal components (PCs). We combine the genotypes and PCs to perform either a score test or a likelihood ratio test to obtain a joint ancestry-adjusted HWE statistic for each variant across all samples.

Although meta-analysis resulted in well-controlled Type I errors for high-coverage data, it was considerably less powerful than RUTH. For example, with moderate excess heterozygosity (θ = -.05) across five ancestries (F_ST_ = .1), RUTH tests identified 20-27% more variants as significant at p < .001 (Figure 2, Table S1) compared to meta-analysis. PRCs also clearly showed better operating characteristics for RUTH and PCAngsd compared to meta-analysis (Figure S2). For example, at an empirical false positive rate of 1%, RUTH showed much greater power (66-68%) than meta-analysis (43%), even though the simulation scenario favors meta-analysis because samples were perfectly classified into distinct subpopulations.

### Application to 1000 Genomes WGS data

Next, we evaluated the performance of various HWE tests in low-coverage (~6x) sequence data from the 1000 Genomes Project. We evaluated three representations of genotypes - (1) raw GT, (2) LD-aware GT, and (3) GL, as described in Materials and Methods. Among chromosome 20 variants, we selected 17,740 high-quality (HQ) variants that are polymorphic in GWAS arrays, and 10,966 low-quality (LQ) variants enriched for genotype discordance in duplicates and trios. Unlike simulation studies, not all LQ variants are necessarily expected to violate HWE, so we consider the proportion of significant LQ variants as a lower bound on the sensitivity to identify significant variants. Similarly, not all HQ variants are necessarily expected to follow HWE, although we expect most to do so, so that the proportion of significant HQ variants serves as an upper bound for the false positive rate.

Consistent with our simulation results, all tests based on raw GTs generated from low-coverage sequence data had severe inflation of false positives (Figure 4A, Table 1). This was true even for HQ variants, presumably due to genotyping errors and bias in raw GTs. Standard HWE tests, which model neither genotype uncertainty nor population structure, showed the highest inflation of false positives at 44% for p < 10^-6^, a threshold commonly used for HWE testing in large genetic studies (Locke *et al*. 2015; Fritsche *et al*. 2016). Modeling population structure substantially reduced inflation, with RUTH tests showing fewer false positives (0.7-1.0% at p < 10^-6^) than meta-analysis (2.0% at p < 10^-6^). False positives were inflated across all methods when using raw GTs.

**Figure 4.**
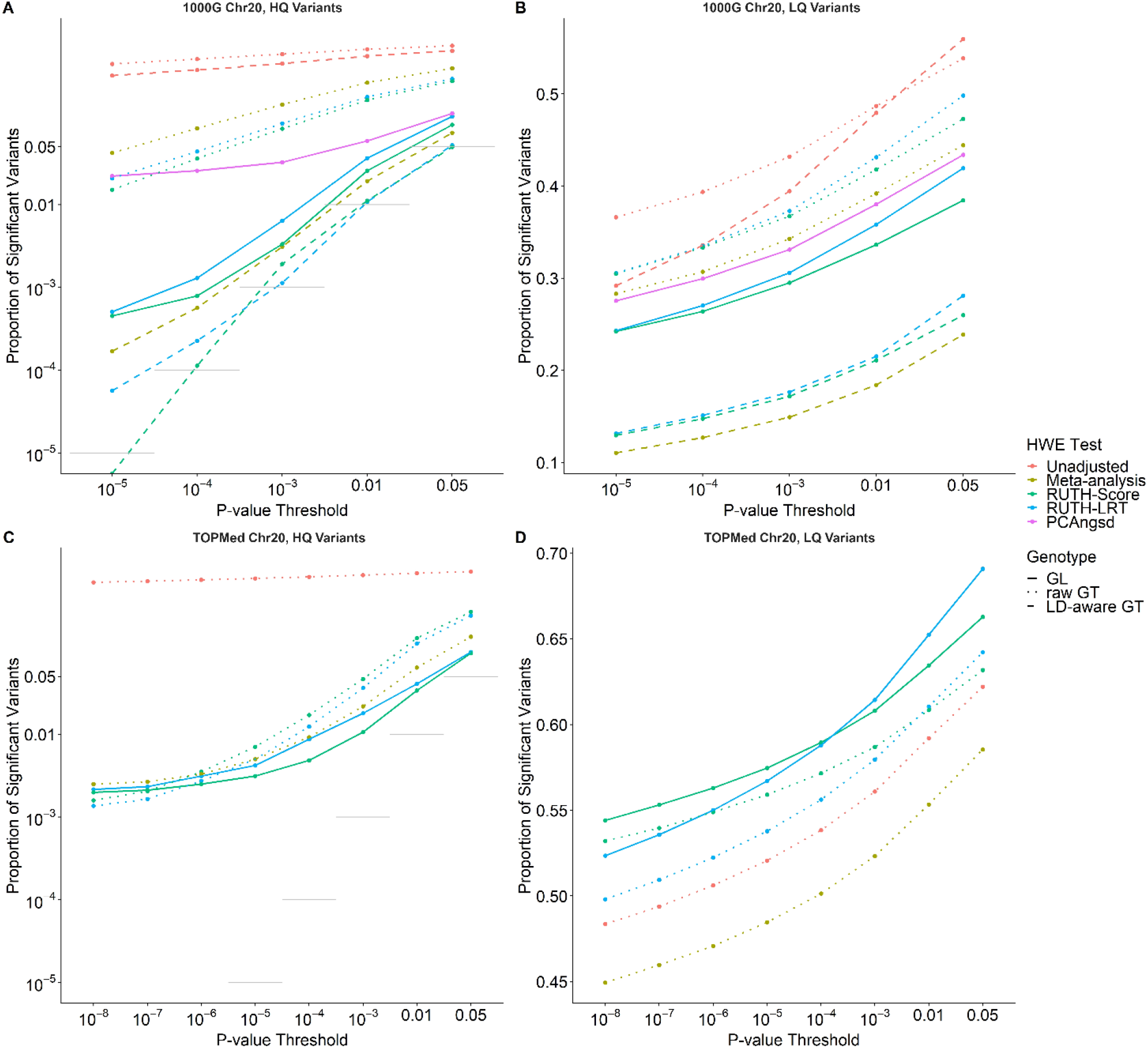
Evaluation of different HWE tests on 1000 Genomes and TOPMed variants. In 1000 Genomes data (A, B), we identified 17,740 “high quality” (HQ) variants and 10,966 “low quality” (LQ) variants in chromosome 20. In TOPMed data (C, D), we identified 17,524 HQ variants and 329,699 LQ variants in chromosome 20. A well-behaved HWE test should maximize the proportion of significant LQ variants while controlling the false positive rate for HQ variants. Dotted gray lines represent targeted Type I error levels if we assume all HQ variants follow HWE. (A) Both the unadjusted test and PCAngsd found substantially more significant variants than expected in the 1000G HQ variant set, while both RUTH and meta-analysis were more conservative. Methods that used raw GTs showed substantial false positive rates, while methods that used GLs and LD-aware GTs had much better control of false positives. (B) In 1000G LQ variants, meta-analysis lagged behind RUTH and the unadjusted test in discovering significant deviation from HWE. RUTH behaved well for HQ variants while having more power to find low-quality variants significantly deviating from HWE. (C) In TOPMed data, the unadjusted test resulted in an excess of false positives. Tests using GL-based genotypes outperformed tests using GT-based genotypes. (D) Methods using GL-based genotypes were able to discover more LQ variants than methods using GT-based genotypes, demonstrating the advantage of accounting for genotype uncertainty in HWE tests.

**Table 1.**
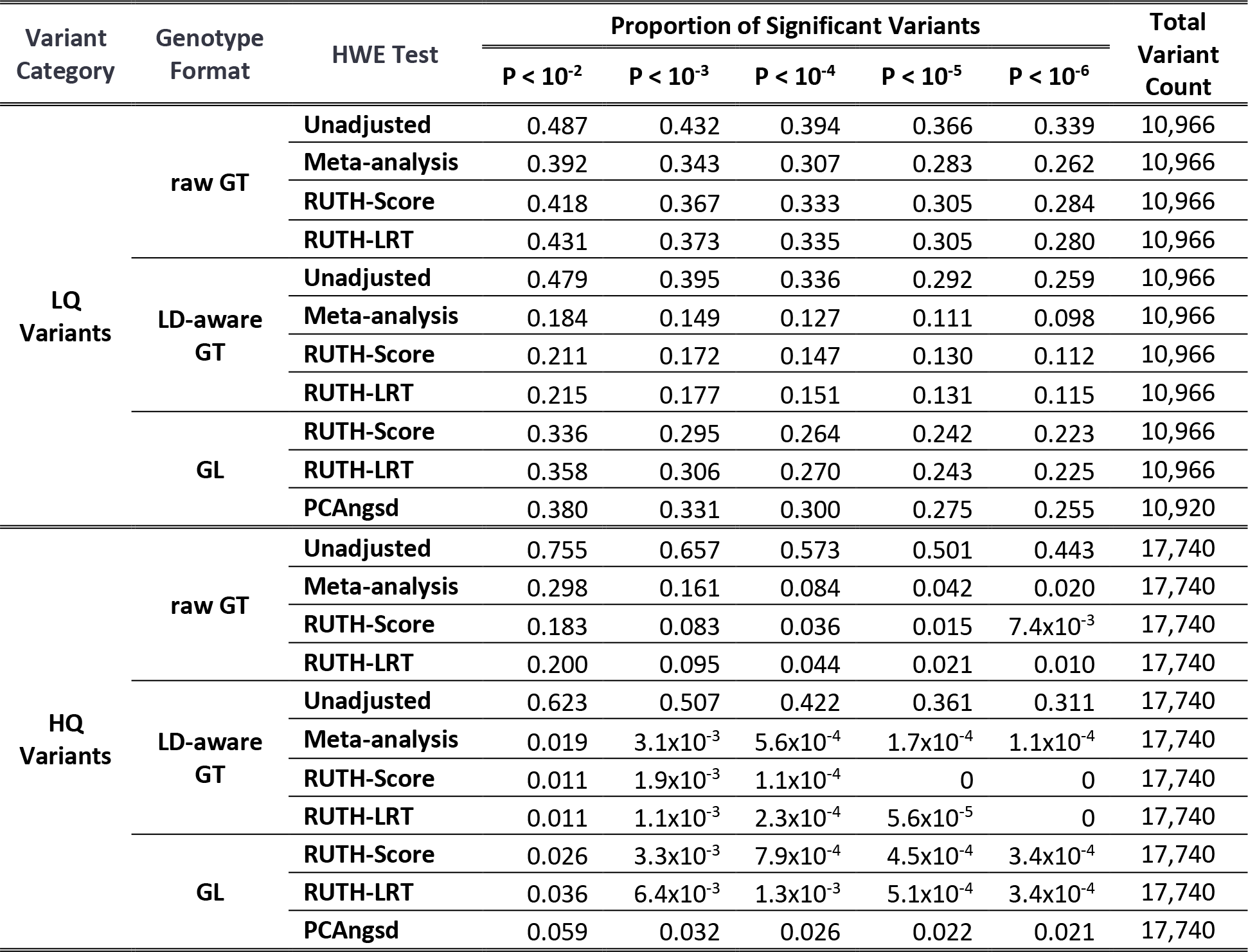
Performance of the unadjusted test, meta-analysis, RUTH, and PCAngsd on 1000 Genomes chromosome 20 variants. The numbers within cells represent the proportions of significant variants under the corresponding testing conditions at the given P-value threshold. We expect our LQ variants to violate HWE at a higher rate than our HQ variants. A well-behaved test is expected to find a high proportion of LQ variants to be significant while maintaining the targeted Type I Error rate in HQ variants. The unadjusted test consistently shows the highest false positive rate among all the tests. HWE tests that rely on raw GTs also show much higher false positive rates than tests that use other genotype representations. RUTH tests were the best at controlling false positives while still maintaining comparable power to the other methods. PCAngsd had a much higher false positive rate than RUTH-based methods, especially at more stringent p-value thresholds.

Consistent with our simulation studies, GL-based RUTH tests reduced false positives even further (0.034% at p < 10^-6^). In contrast to our simulations, PCAngsd demonstrated considerably higher false positives than RUTH (2.1% at p < 10^-6^), likely because PCAngsd estimates PCs from the input data without the ability to use externally provided PCs (see Discussion). The sensitivity for detecting significant LQ variants was also consistent with our simulations (Figure 4B, Table 1). GL-based tests, which showed better control of false positives, identified 22-25% of LQ variants as significant at p < 10^-6^.

Strikingly, while using LD-aware GTs reduced false positives with adjusted tests, it was at the expense of substantially reduced sensitivity to detect LQ variants. The false positive rates of any adjusted test with LD-aware GTs were uniformly lower than those of any GL- and raw GT-based tests across all p-value thresholds (Figure 4A). However, sensitivity was also substantially reduced with LD-aware genotypes (Figure 4B). For example, at p < 10^-6^, GL-based RUTH tests identified 22-23% of LQ variants significant, while using LD-aware GTs halved the proportions. Running meta-analysis with LD-aware GTs reduced sensitivity even further, likely because the implicit HWE assumption in the LD-aware genotype refinement algorithms may have further reduced false positives and sensitivity by altering the LD-aware genotypes to conform to HWE.

We evaluated PRCs between HQ and LQ variants to further evaluate this tradeoff. The results clearly demonstrated that HWE tests using LD-aware GTs are substantially less robust than tests on other genotype representations (Table S2, Figure S3A). For example, for the RUTH score test, when LD-aware GTs identified 0.1% of HQ variants as significant, 17% of LQ variants were identified as significant. However, with raw GT and GL, 24~27% were identified as significant at the same threshold. Even fewer were significant in meta-analysis with LD-aware GTs (13%). Similar trends were observed across all thresholds, suggesting that using LD-aware GTs results in substantially poorer operating characteristics than other genotype representations. As more accurate genotyping in LD-aware genotype refinement is expected to improve the performance of QC metrics compared to raw GTs, these results are quite striking, and highlight a potential oversight in using LD-aware genotypes in various QC metrics for sequence-based genotypes.

### Application to TOPMed Deep WGS data

We evaluated the various HWE tests on a subset of the Freeze 5 variant calls from the high-coverage (~37×) whole genome sequence (WGS) data in the TOPMed Project (Taliun *et al*. 2019). We identified 17,524 HQ variants and 329,699 LQ variants using the same criteria used for 1000G variants and evaluated raw GTs and GLs. We did not evaluate PCAngsd due to excessive computational time (see “Computational cost” below).

We first evaluated the false positive rates of different HWE tests indirectly by using HQ variants. With a >20-fold larger sample size than 1000G, we identified more significant HQ variants, while the false positive rates were still reasonable with adjusted tests. At p < 10^-6^, 74% of HQ variants were significant with unadjusted tests, while the adjusted GL-based tests identified ~0.3% at p < 10^-6^ (Figure 4C-D, Table 2). Adjusted GT-based tests had only slightly higher levels of false positives at p < 10^-6^. However, inflation was more noticeable at less stringent p-value thresholds suggesting that GL-based tests may be needed for larger sample sizes.

**Table 2.** Performance of the unadjusted test, meta-analysis, and RUTH on TOPMed freeze 5 chromosome 20 variants. The numbers within cells represent the proportions of significant variants under the corresponding testing conditions at the given P-value threshold. These results are based on tests that used likelihood-based genotype representations as input. A well-behaved test should reduce the number of significant high-quality (HQ) variants while increasing the number of significant low-quality (LQ) variants. The unadjusted test had a greatly inflated false positive rate for HQ variants while showing a lower true positive rate for LQ variants. While meta-analysis performed better for HQ variants, it had reduced power to find LQ variants to be significant. RUTH performed the best, with fewer false positives (significant HQ variants) compared to both the unadjusted test and meta-analysis, while at the same time finding more true positives (significant LQ variants).

| Variant set | Genotype Format | HWE Test | Proportion of Significant Variants |  |  |  |  | Total Variant Count |
| --- | --- | --- | --- | --- | --- | --- | --- | --- |
| | | | $P < 10^{-2}$ | $P < 10^{-3}$ | $P < 10^{-4}$ | $P < 10^{-5}$ | $P < 10^{-6}$ | |
| LQ Variants | raw GT | Unadjusted | 0.592 | 0.561 | 0.539 | 0.521 | 0.506 | 329,699 |
|  | raw GT | Meta-analysis | 0.554 | 0.524 | 0.502 | 0.485 | 0.471 | 329,699 |
|  | raw GT | RUTH-Score | 0.608 | 0.587 | 0.572 | 0.559 | 0.549 | 329,699 |
|  | GL | RUTH-Score | 0.635 | 0.608 | 0.590 | 0.575 | 0.563 | 329,699 |
|  | raw GT | RUTH-LRT | 0.610 | 0.580 | 0.556 | 0.538 | 0.522 | 329,699 |
|  | GL | RUTH-LRT | 0.653 | 0.615 | 0.588 | 0.567 | 0.550 | 329,699 |
| HQ Variants | raw GT | Unadjusted | 0.890 | 0.842 | 0.800 | 0.766 | 0.736 | 17,524 |
| | raw GT | Meta-analysis | 0.065 | 0.022 | $9.0 \times 10^{-3}$ | $4.8 \times 10^{-3}$ | $3.3 \times 10^{-3}$ | 17,524 |
| | raw GT | RUTH-Score | 0.145 | 0.047 | 0.172 | $7.1 \times 10^{-3}$ | $3.5 \times 10^{-3}$ | 17,524 |
| | GL | RUTH-Score | 0.034 | 0.011 | $4.9 \times 10^{-3}$ | $3.1 \times 10^{-3}$ | $2.5 \times 10^{-3}$ | 17,524 |
| | raw GT | RUTH-LRT | 0.125 | 0.036 | 0.012 | $5.0 \times 10^{-3}$ | $2.7 \times 10^{-3}$ | 17,524 |
| | GL | RUTH-LRT | 0.041 | 0.018 | $8.5 \times 10^{-3}$ | $4.3 \times 10^{-3}$ | $3.1 \times 10^{-3}$ | 17,524 |

Next, we evaluated the proportions of LQ variants found to be significant by different tests to indirectly evaluate their statistical power. GT- and GL-based RUTH tests showed similar power, while meta-analysis showed considerably lower power. For example, at p < 10^-6^, meta-analysis identified 47% of LQ variants as significant, while RUTH tests identified 54-58%. This pattern was similar across different p-value thresholds (Figure 4C-D) or choices of LQ variants (Table S3, Figure S4). Our results suggest that GL-based RUTH tests are suitable for testing HWE for tens of thousands of deeply sequenced genomes with diverse ancestries, but that using raw GTs will also result in a comparable performance at typically used HWE p-value thresholds (e.g. p < 10^-6^) when performing QC without access to GLs.

We used PRCs to evaluate the tradeoff between empirical false positive rates and power. Consistent with previous results, the GL-based RUTH test showed the best tradeoff between false positives and power, while the GT-based RUTH test and meta-analysis were slightly less robust but largely comparable (Figure S3). Notably, when we evaluated the different methods at an empirical false positive rate of 0.1%, RUTH score tests had ~4% higher power than RUTH LRT for both raw GTs and GLs (Figure S5–6).

### Impact of ancestry estimation accuracy on HWE tests

So far, our evaluations relied on genetic ancestry estimates carefully determined with sophisticated methods (see Materials and Methods). However, simpler approaches may be used instead during the variant QC step, which may affect the performance of adjusted HWE tests. We evaluated whether the number of PC coordinates affected the performance of RUTH tests by comparing the performance of RUTH tests when using 2 PCs to using 4 PCs (default). The results from both simulated and real datasets consistently demonstrated that using 4 PCs led to substantially reduced Type I errors compared to using 2 PCs at a similar level of power (Table S2, Table S4, Figure S7). PRCs also clearly showed that using 4 PCs was more robust against population structure across both simulated and real datasets (Figure S8).

We also evaluated whether the classification accuracy of subpopulations affected the performance of meta-analysis. Instead of assigning 1000 Genomes individuals into five continental populations, we used the k-means algorithm on those samples’ top 2 PCs to classify them into 3 crude subpopulations (Figure S9). This led to a much higher false positive rate with virtually no increase in true positives (Figure S10, Table S2). We saw the same pattern in simulated data (Figure S8, Table S5).

### Computational cost

We compared the computational costs of RUTH and PCAngsd for simulated and real data. RUTH has linear time complexity to sample size, while PCAngsd appears to have quadratic time complexity (Tables 3, S6). RUTH also has low memory requirement compared to PCAngsd (for example, 14 MB vs 2 GB for 1000 Genomes data). Extrapolating our results to the whole genome scale, analyzing 1000 Genomes (i.e. 80 million variants) is expected to take 120 CPU-hours for RUTH, and 3,200 CPU-hours for PCAngsd (with >1 TB memory consumption). Additionally, RUTH can be parallelized into smaller regions in a straightforward manner.

**Table 3.** Runtimes for RUTH and PCAngsd on simulated data. We simulated 10,000 genotype likelihood-based variants for varying numbers of samples. Wall time indicates total runtime, while user time is the amount of time the CPUs spent running each program. All programs were run in single-threaded mode. System processes make up the difference between the two values, with a majority consisting of file I/O. We used VCF files with GL fields in RUTH and converted them to Beagle3 format for PCAngsd. The RUTH likelihood ratio test (LRT) was the fastest method, with the score test about 60% slower. PCAngsd was about 10 times slower than RUTH-LRT with the smallest sample sizes and over 400 times slower with our largest tested size of 50,000 samples.

| Sample Size | Wall Time (s) |  |  | User Time (s) |  |  |
| --- | --- | --- | --- | --- | --- | --- |
|  | RUTH-LRT | RUTH-Score | PCAngsd | RUTH-LRT | RUTH-Score | PCAngsd |
| **1,000** | 16.21 | 27.24 | 173.11 | 16.16 | 27.09 | 172.37 |
| **2,000** | 32.19 | 54.63 | 347.10 | 31.94 | 54.51 | 345.58 |
| **5,000** | 82.80 | 136.44 | 1,124.83 | 81.81 | 136.20 | 1,102.85 |
| **10,000** | 165.48 | 273.67 | 7,396.00 | 163.88 | 273.27 | 7,235.91 |
| **20,000** | 336.75 | 553.92 | 38,807.67 | 332.06 | 553.05 | 37,338.69 |
| **50,000** | 902.81 | 1,438.32 | 461,971.33 | 886.67 | 1,435.87 | 403,296.5 |

## DISCUSSION

RUTH is a unified, flexible, and robust approach to incorporate genetic ancestry and genotype uncertainty for testing Hardy-Weinberg Equilibrium capable of handling large amounts of genotype data with structured populations. Sha and Zhang (2011) proposed HWES, an HWE test for structured populations, to address some of these challenges, but it has not been widely used due to the lack of an implementation that supports widely used genotype data formats (e.g. PED, BED, VCF, or BCF) and inability to handle imputed or uncertain genotypes. Hao and colleagues (2016) proposed sHWE which can only handle best-guess (hard call) genotypes (i.e. 0, 1, or 2 for biallelic variants) and does not account for genotype uncertainty. Meisner and Albrechtsen (2019) proposed PCAngsd to address some of these issues, but it does not support the standard VCF/BCF formats for sequence-based genotypes, and its current implementation scales poorly with genome-wide analyses of large samples.

Similar to previous studies (Sha and Zhang 2011; Hao *et al*. 2016), our proposed framework uses individual-specific allele frequencies rather than allele frequencies pooled across all samples to systematically account for population structure in HWE tests. Unlike previous studies, we model genotype uncertainty in sequence-based genotypes in a likelihood-based framework. We implemented two RUTH tests – a score test and a likelihood ratio test (LRT) – to test for HWE under population structure for genotypes with uncertainty. While RUTH LRT is similar to the independently developed PCAngsd, the software implementation of RUTH is more flexible, scales much better to large studies, and supports the standard VCF format.

We provide a comprehensive evaluation of various approaches for testing HWE using simulated and real data. Our results demonstrated that modeling population stratification is necessary for HWE tests on heterogenous populations. We showed that accounting for genotype uncertainty via genotype likelihoods performs substantially better than testing HWE with best-guess genotypes, especially for low-coverage sequenced genomes. Importantly, we included the evaluations for an unpublished but commonly used approach – meta-analysis across stratified subpopulations, cohorts, or batches. Our results demonstrate that meta-analysis may be effective in reducing false positives, but at the expense of substantially reduced power compared to RUTH.

We observed that the current implementation of PCAngsd does not scale well to large-scale sequencing data, though in principle it can be implemented more efficiently, because the underlying HWE test itself is similar to RUTH LRT. PCAngsd requires loading all genotypes into memory, which is often infeasible for large sequencing studies. For example, loading all of 1000 Genomes will require ~4.8 TB of memory. In our evaluation of 1000G chromosome 20 variants, the inability of PCAngsd to estimate PCs from the whole genome may have contributed to the observed difference in results from RUTH compared to our simulation studies.

Although our 1000G experiments demonstrated the unexpected result that using raw GTs had better sensitivity than using LD-aware GTs at the same empirical false positive rates for low-coverage data, we do not advocate using raw GTs for low-coverage sequence data. First, the results for raw GTs were still consistently less robust than GL-based RUTH tests. Moreover, it would be tricky to determine an appropriate p-value threshold when the false positives are severely inflated. Therefore, we strongly advocate using GL-based RUTH tests for robust HWE tests with low-coverage sequence data. For the now more typical high-coverage sequence data, GL-based tests are still preferred, but GT-based RUTH tests should be acceptable for cases in which genotype likelihoods are unavailable.

Our experiment compared using 2 vs 4 PCs only because *verifyBamID2* software tool estimated up to 4 PCs projected onto HGDP panel by default (Zhang *et al*. 2020). Because our method focuses on testing HWE during the QC steps in sequence-based variant calls, a curated version of PCs, estimated from sequenced cohort themselves, may not be readily available at the time of HWE test. However, it is possible to use a larger number of PCs (e.g. >10 PCs) if available at the time of HWE test. We expect that a larger number of PCs will account for finer-grained population structure and may benefit the performance of HWE test, but additional experiments are needed to quantify the impact of using larger number of PCs.

Our results demonstrate that RUTH score and LRT tests perform similarly in simulated and experimental datasets. Overall, the RUTH-LRT was slightly more powerful than the RUTH-score test at the expense of slightly greater false positive rates, although this tendency was not consistent. We observed that the RUTH tests tended to be slightly more powerful in identifying deviation from HWE in the direction of excess heterozygosity than excess homozygosity when compared to adjusted meta-analysis. These results might be caused by the difference between our model-based asymptotic tests compared to the exact test used in meta-analysis.

We did not evaluate our methods on imputed genotypes in this manuscript. Because imputed genotypes implicitly assume HWE, we suspect that HWE tests based on imputed genotypes may have reduced power compared to directly genotyped variants. It is possible to use approximate genotype likelihoods instead of best-guess genotypes for imputed genotypes, but this requires genotype probabilities, not just the genotype dosages. If genotype probabilities Pr (*g_i_* = *G*|*Data_i_*) are available, they can be converted to genotype likelihoods 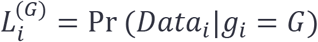 using Bayes’ rule by modeling Pr(*g_i_* = *G*) as a binomial distribution based on allele frequencies (which implicitly assumes HWE). However, similar to LD-aware genotypes in low-coverage sequencing, the power of HWE tests with imputed genotypes may be poor. Further evaluation is needed to understand how useful this approximation will be compared to alternative methods including the use of best-guess imputed genotypes.

Our methods have room for further improvement. First, we used a truncated linear model for individual-specific allele frequencies for computational efficiency. Although such an approximation was demonstrated to be effective in practice (Zhang *et al*. 2020), applying a logistic model or some other more sophisticated model may be more effective in improving the precision and recall of RUTH tests. Second, we did not attempt to model or evaluate the effect of admixture in our method. Because HWE is reached in two generations with random mating, accounting for admixed individuals may only have marginal impact. On the other hand, admixture can lead to higher observed heterozygosity. It may be possible to improve RUTH by explicitly modeling and adjusting for the effect of admixture on individual-specific allele frequencies. Systematic evaluations focusing on admixed populations are needed to evaluate RUTH’s performance on such samples, and whether an admixture adjustment is necessary. Third, RUTH tests do not account for family structure. We suspect that the apparent inflation of Type I error for the TOPMed data was partially due to sample relatedness. Accounting for family structure in other ways, for example using variance components models, will require much longer computational times and may not be feasible for large-scale datasets. Fourth, RUTH currently does not directly support imputed genotypes or genotype dosages. In principle, it is possible to convert posterior probabilities for imputed genotypes into genotype likelihoods to account for genotype uncertainty (by using individual-specific allele frequencies). However, because most genotype imputation methods implicitly assume HWE, we suspect that HWE tests on imputed genotypes will be underpowered, similar to our observations with LD-aware genotypes in the 1000 Genomes dataset, even though explicitly modeling posterior probabilities may slightly mitigate this reduction in power.

In summary, we have developed and implemented robust and rapid methods and software tools to enable HWE tests that account for population structure and genotype uncertainty. We performed comprehensive evaluations of both our methods and alternative approaches. Our tools can be used to evaluate variant quality in very large-scale genetic data sets, with the ability to handle standard VCF formats for storing sequence-based genotypes. Our software tools are publicly available at http://github.com/statgen/ruth.

## Acknowledgements

This work was supported by NIH grants HL137182 (from NHLBI), HG009976 (from NHGRI), HG007022 (from NHGRI), DA037904 (from NIDA), HL117626-05-S2 (from NHLBI), and MH105653 (from NIMH). Molecular data for the Trans-Omics in Precision Medicine (TOPMed) program was supported by the National Heart, Lung and Blood Institute (NHLBI). Core support including centralized genomic read mapping and genotype calling, along with variant quality metrics and filtering were provided by the TOPMed Informatics Research Center (3R01HL-117626-02S1; contract HHSN268201800002I). Core support including phenotype harmonization, data management, sample-identity QC, and general program coordination were provided by the TOPMed Data Coordinating Center (R01HL-120393; U01HL-120393; contract HHSN268201800001I). We gratefully acknowledge the studies and participants who provided biological samples and data for TOPMed.

TOPMed source studies and sample counts are described in Table S7. Acknowledgements for TOPMed omics support are detailed in Table S8. Full TOPMed study acknowledgements are listed in Supplementary File S1.

## List of Supplementary Figures, Tables, and File

**Figure S1.**
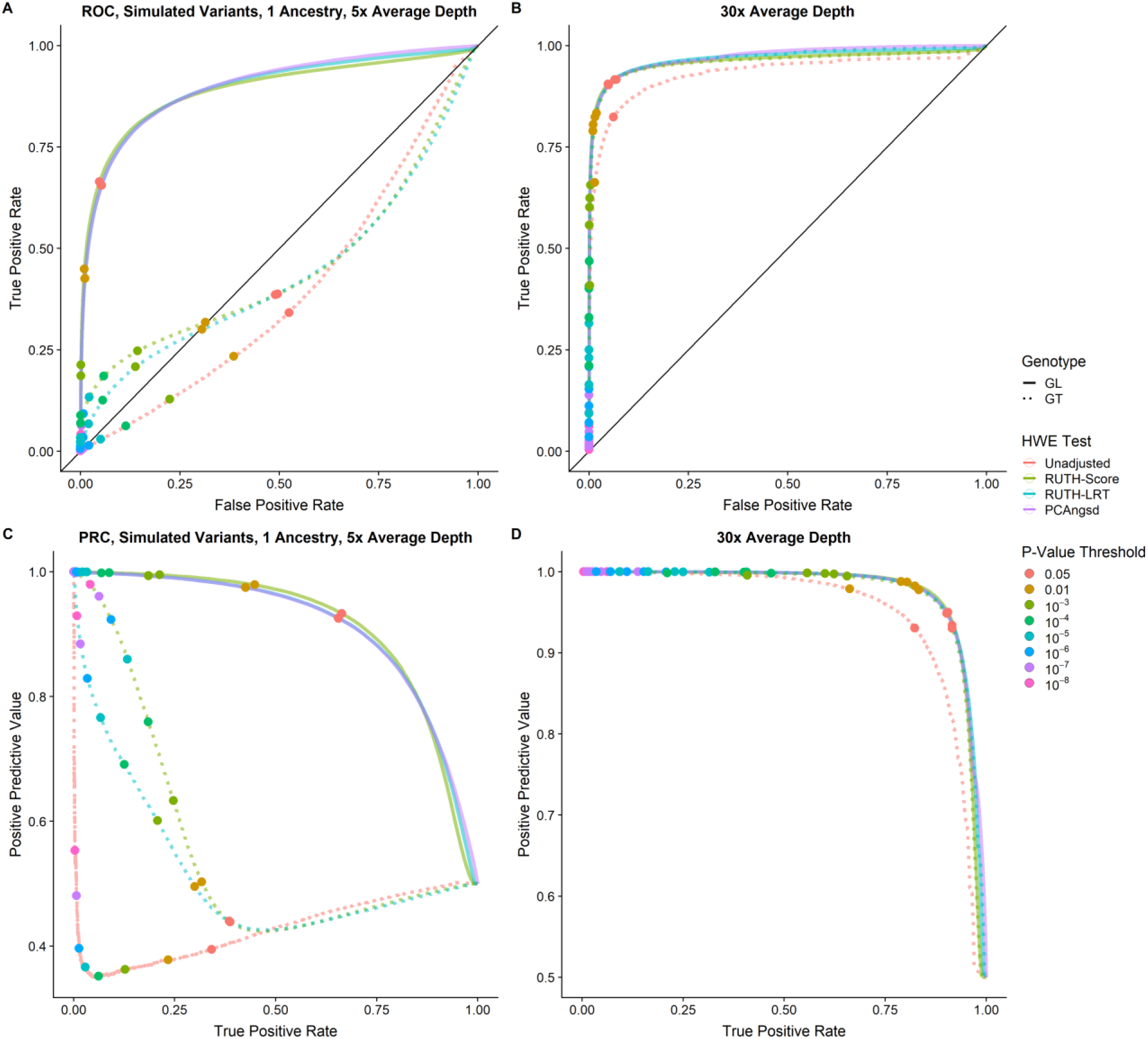
ROC and PRC for simulated single-ancestry data. For both low coverage (A, C) and high coverage (B, D) settings, 500,000 variants were generated from 5,000 samples arising from a single ancestry, with half of the variants as true positives (θ = −0.05) and half of the variants as true negatives (θ = 0). The colors of the lines correspond to the different HWE tests, while the colors of the points correspond to different P-value thresholds. In all cases, the unadjusted test performed the worst. For low-coverage data, tests using GT-based genotypes performed poorly due to their inability to capture the effects of genotype uncertainty, whereas tests using GL-based genotypes performed much better. The difference was negligible in high-coverage genotype data.

**Figure S2.**
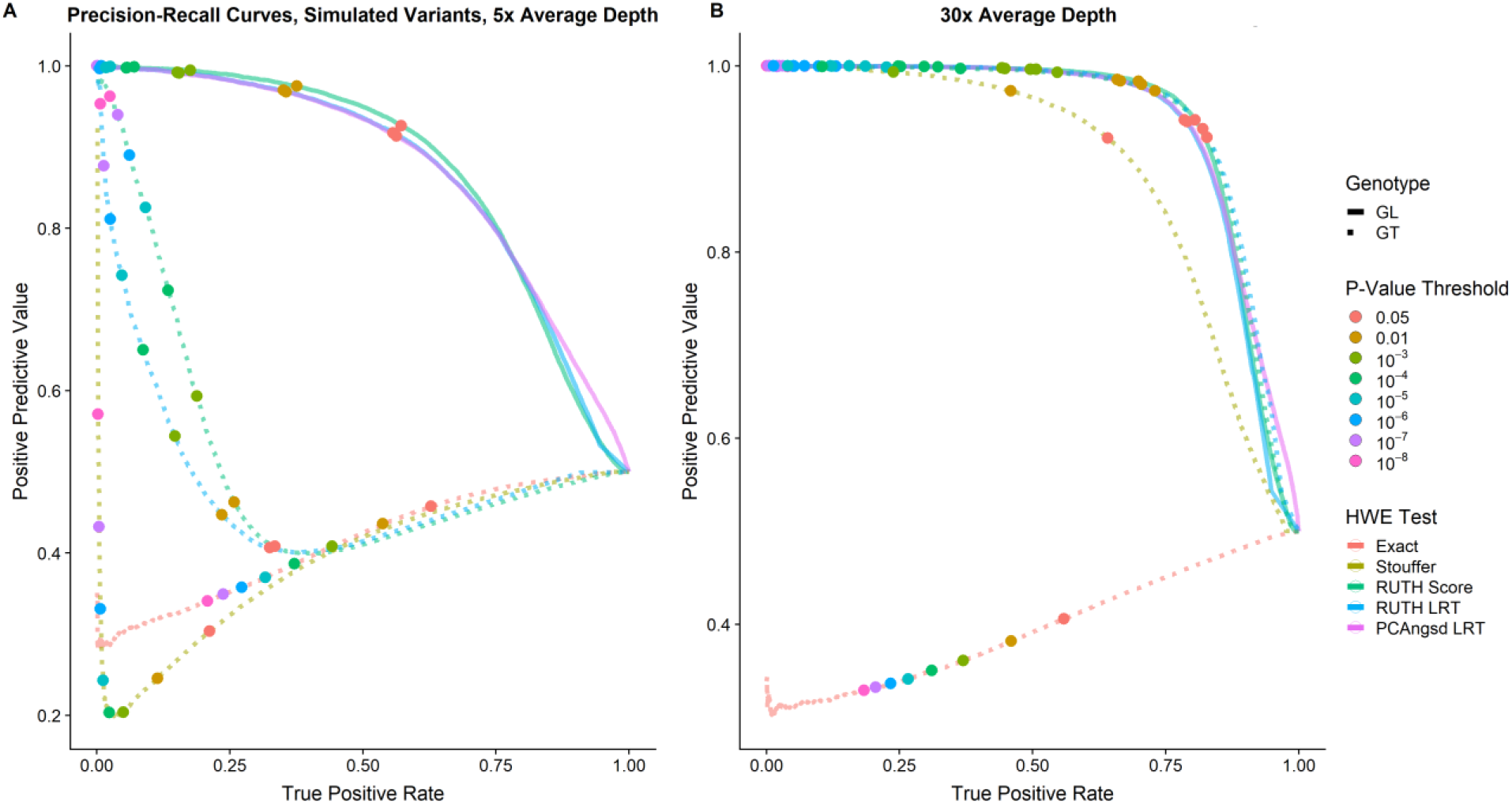
Precision-recall curves for simulated data with multiple ancestries. We generated Precision-recall curves to evaluate the tradeoff between the different HWE tests’ ability to identify true positive variants while minimizing the misidentification of true negative variants as significantly departing from HWE. We analyzed 50,000 true positive and 50,000 true negative variants in 5,000 samples arising from 5 different ancestries with an average simulated depth of (A) 5x and (B) 30x. True negative variants are defined as variants with the HWE deviation parameter θ = 0. True positives are defined as variants with θ = −0.05. The True Positive Rate (TPR) is defined to be the proportion of variants with θ = −0.05 that are significant at a given P-value threshold, while the Positive Predictive Value (PPV) is defined as the proportion of significant variants with θ = −0.05 at the same P-value threshold. Selected p-value thresholds are indicated with colored circles. For low-depth genotypes, in the presence of high genotype uncertainty, GL-based HWE tests performed relatively well, while GT-based tests performed poorly. For high-depth genotypes, with low genotype uncertainty, all methods adjusting for population structure performed relatively well.

**Figure S3.**
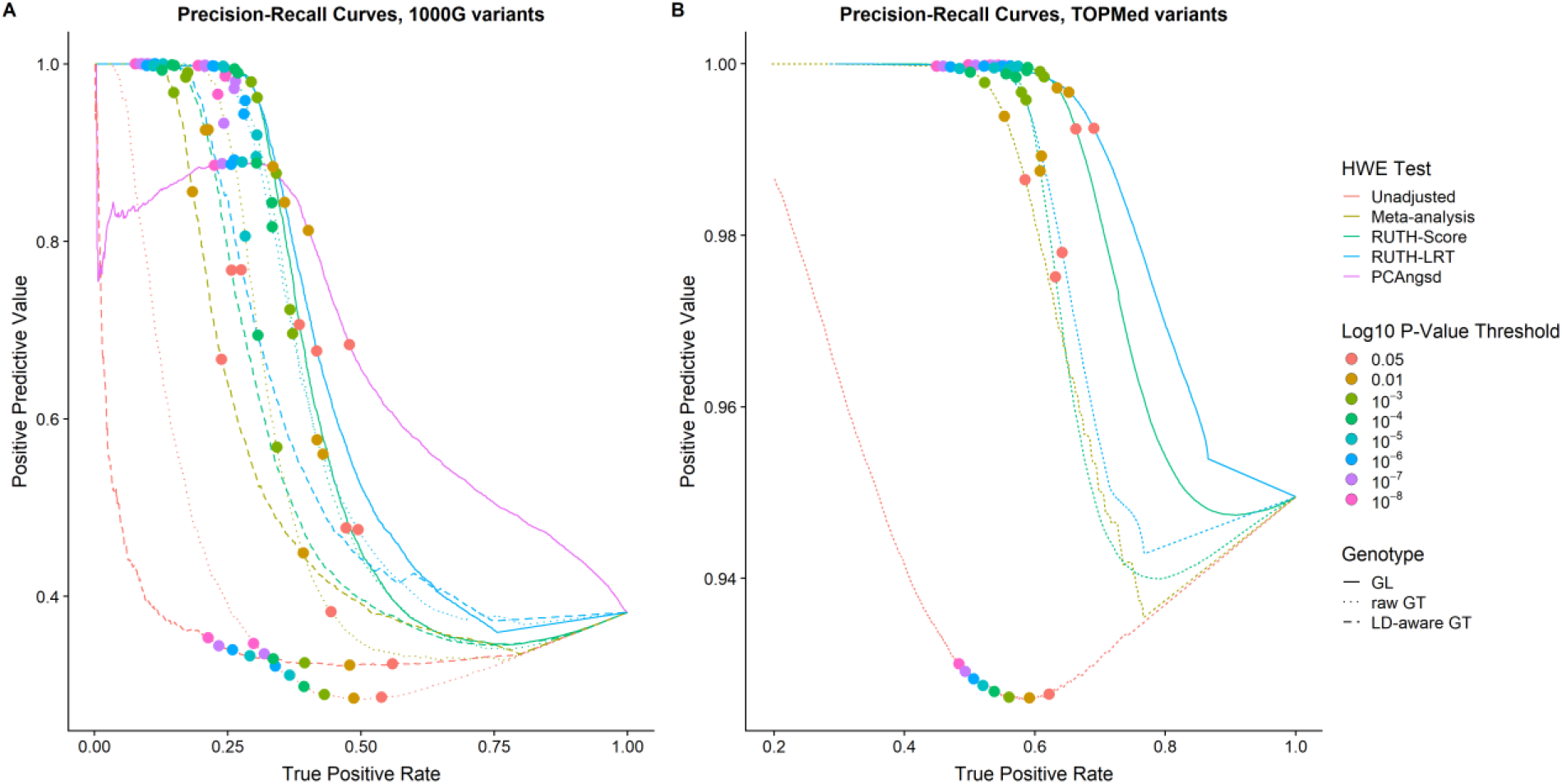
Precision-recall curves for 1000G and TOPMed variants. We defined positive variants as those with a high level of Mendelian inconsistency in family-based TOPMed data, and negative variants as those found in the intersection of the Illumina Omni2.5 and HapMap3 variant site lists. (A) For low-coverage sequence data found in 1000G, tests using GL-based genotypes (solid lines) generally performed better than tests using any GT-based genotypes (dotted and dashed lines). Both the unadjusted test and meta-analysis performed much worse than all other methods. (B) For high-coverage sequence data found in TOPMed, tests using GL-based genotypes retained their improved performance over tests using GT-based genotypes.

**Figure S4.**
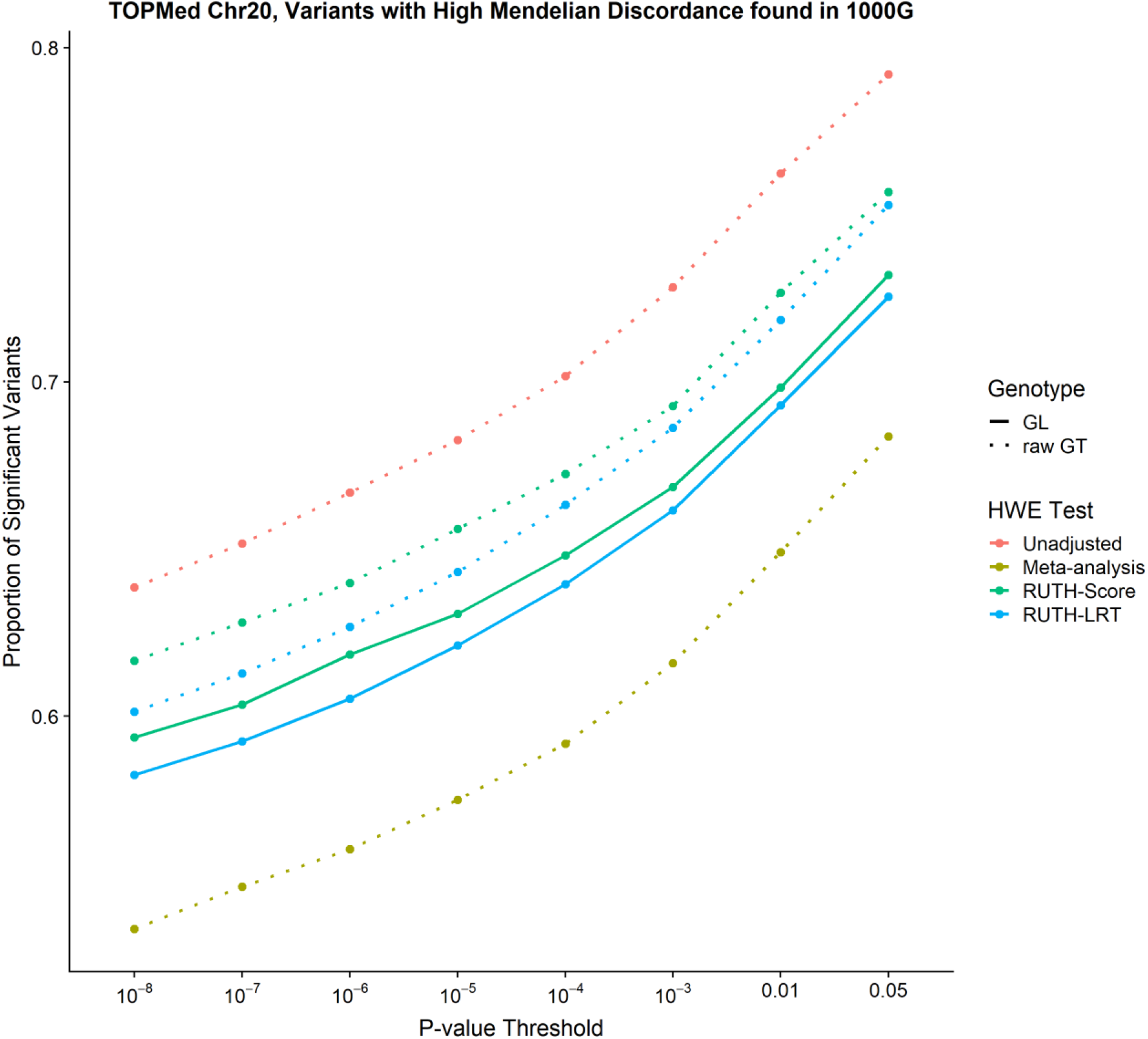
Results of testing TOPMed variants found in 1000G variant list. This analysis contains 10,966 TOPMed variants found to be discordant in TOPMed family data and overlapping with 1000G discordant variants, as opposed to all 329,699 discordant TOPMed variants (as seen in Figure 4D). Our results are similar to those for 1000G discordant variants (Figure 4B), suggesting that the differences between the patterns observed in 1000G and TOPMed results may have been caused by the difference in allele frequency distributions in the two data sets (Table S1).

**Figure S5.**
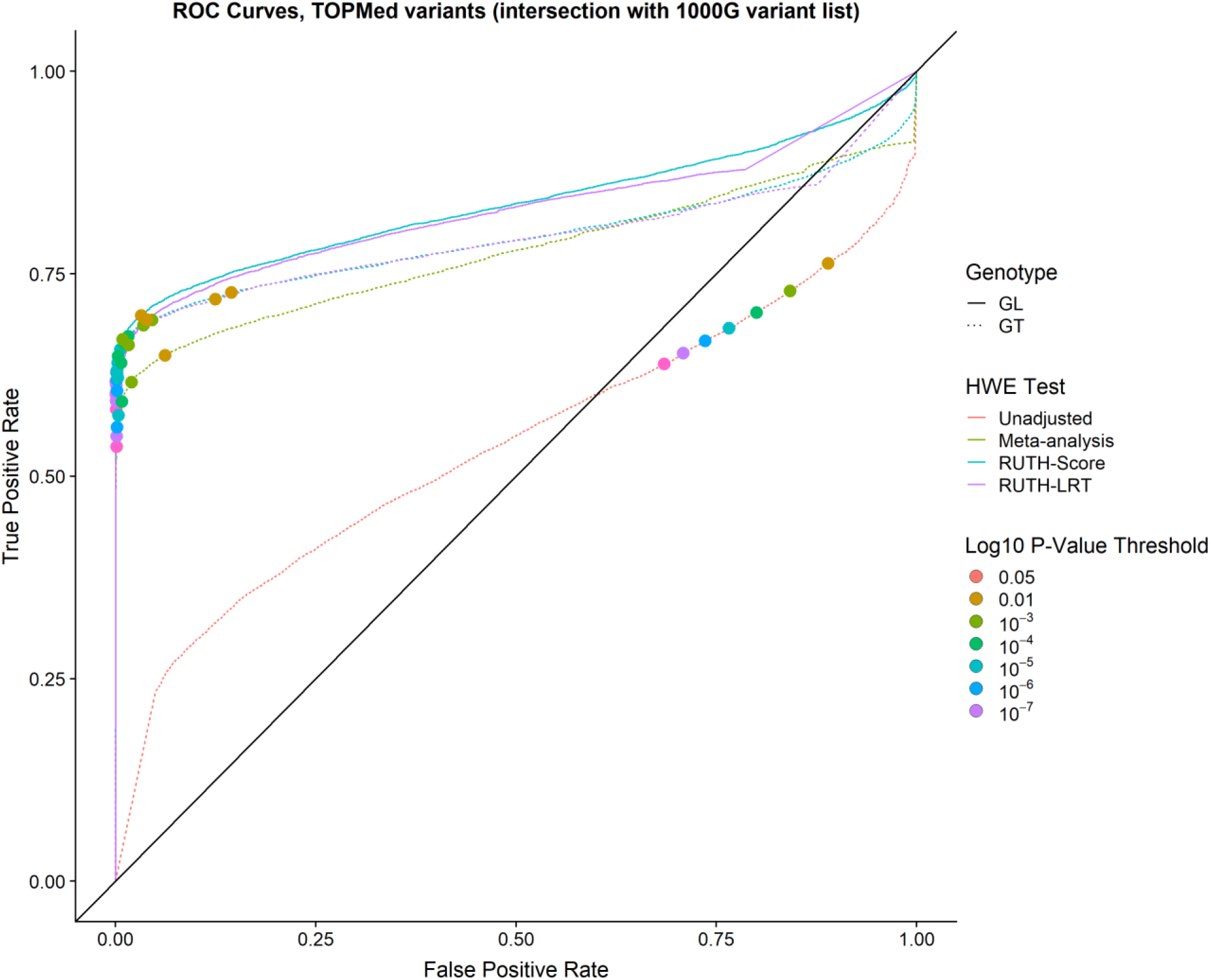
ROC curves for TOPMed variants found in 1000G variant list. GL-based tests have the best overall performance among the different methods.

**Figure S6.**
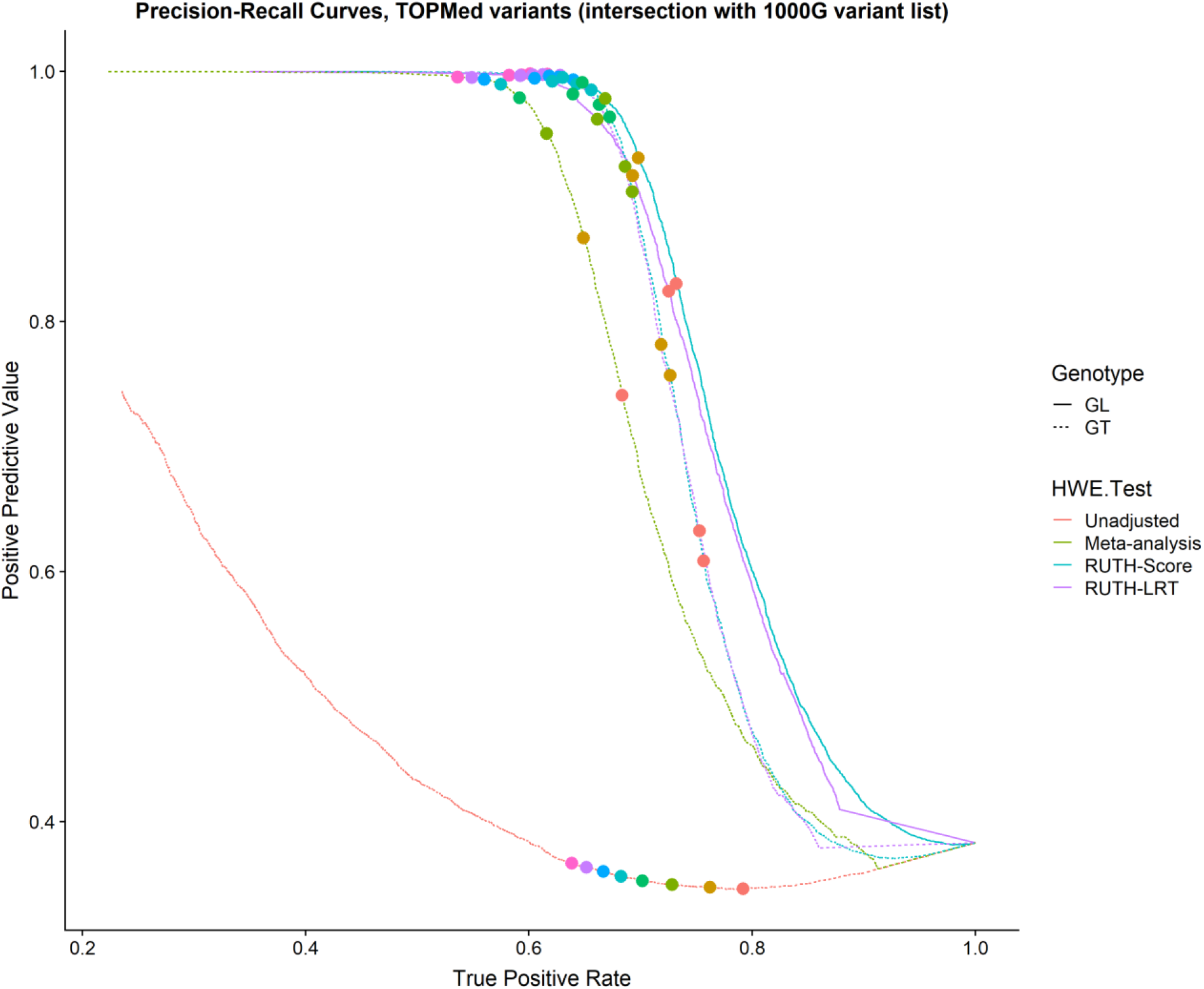
PRC curves for TOPMed variants found in 1000G variant list. RUTH tests using GLs offer the best balance between finding true positives and maximizing positive predictive value.

**Figure S7.**
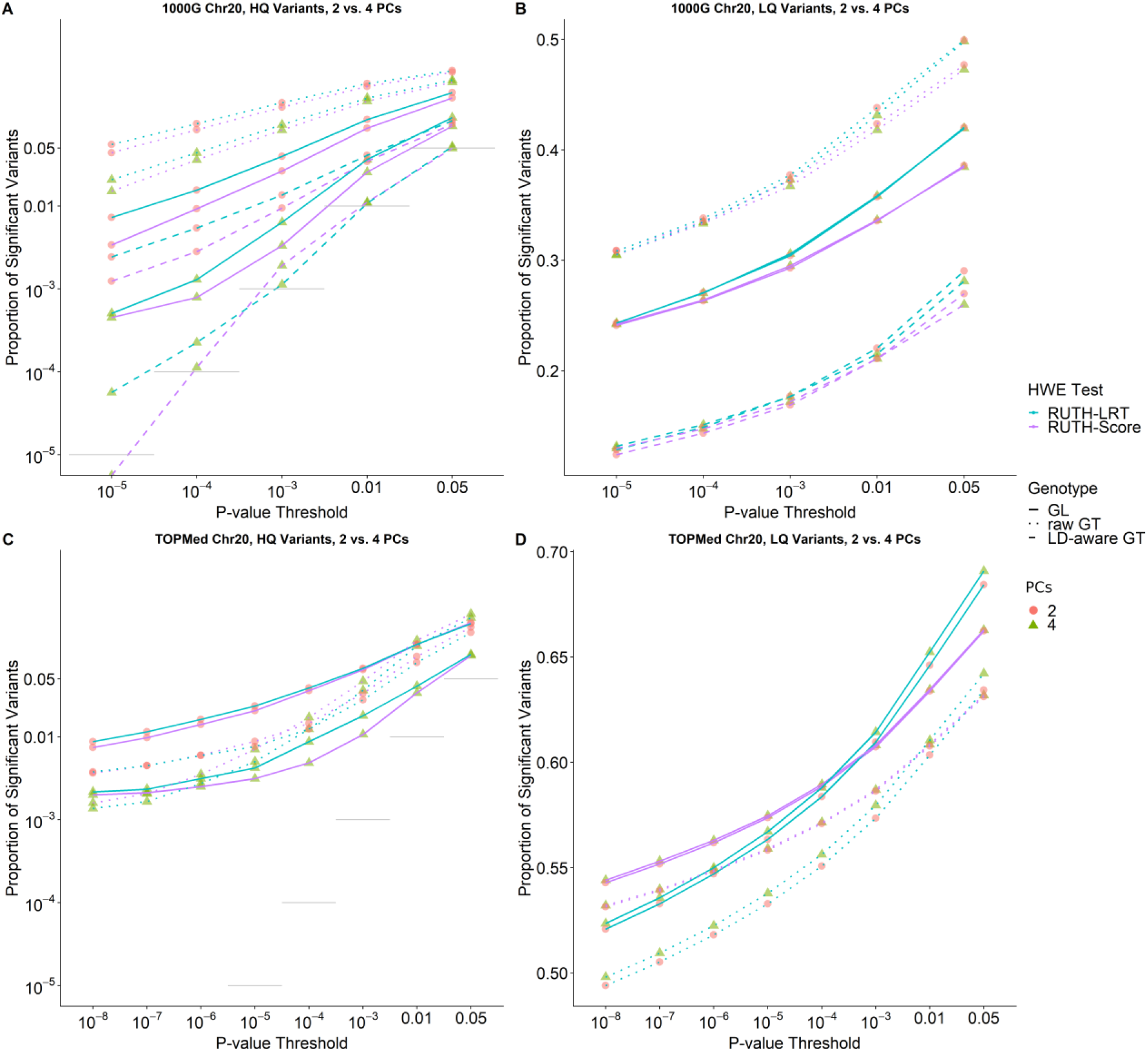
Results of testing 1000G and TOPMed variants with RUTH using two vs. four PCs. Using only 2 PCs lead to noticeably worse performance, especially for GL-based tests. (A) In 1000 Genomes data, using only 2 PCs leads to much higher false positives in HQ variants for both RUTH-Score and RUTH-LRT compared to using 4 PCs. (B) Tests on LQ variants with 2 PCs appear to have modestly higher power than tests using 4 PCs, but this is mainly due to the much higher false positive rate. (C) For HQ variants in TOPMed, tests using only 2 PCs have substantially higher false positive rate than tests using 4 PCs for GL-based tests, while GT-based tests are comparable. (D) Surprisingly, GL-based tests using 4 PCs discovered more significant LQ variants compared to GL-based tests using 2 PCs, even though GL-based tests using 2 PCs had a higher false positive rate in HQ variants.

**Figure S8.**
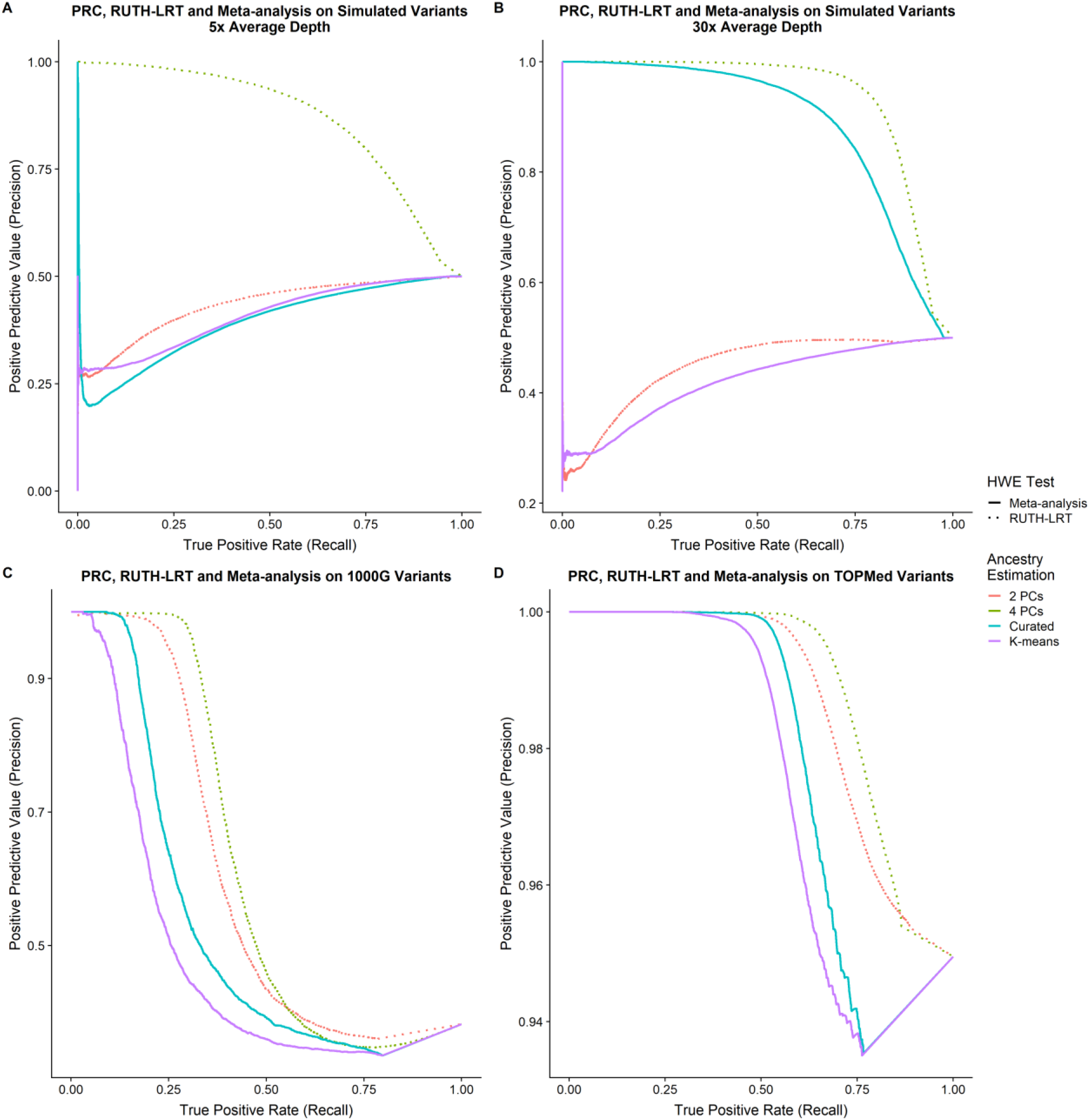
Effect of ancestry estimation accuracy on Precision-Recall Curves. We evaluated the effect of using 2 vs. 4 principal components on the performance of RUTH-LRT, and the effect of using our nearest-neighbor algorithm (“curated”) vs. k-means for subpopulation classification of samples on the performance of meta-analysis on (A) low-depth simulated data, (B) high-depth simulated data, (C) 1000G variants, and (D) TOPMed variants. We simulated null variants with θ = 0 and alternative variants with θ = −0.05, with a fixation index of 0.1 for 5,000 samples from 5 ancestries (1,000 samples each). RUTH-LRT used GL-based genotypes, and meta-analysis used raw GT-based genotypes. K-means classification for simulated data was performed assuming 3 subpopulation clusters.

**Figure S9.**
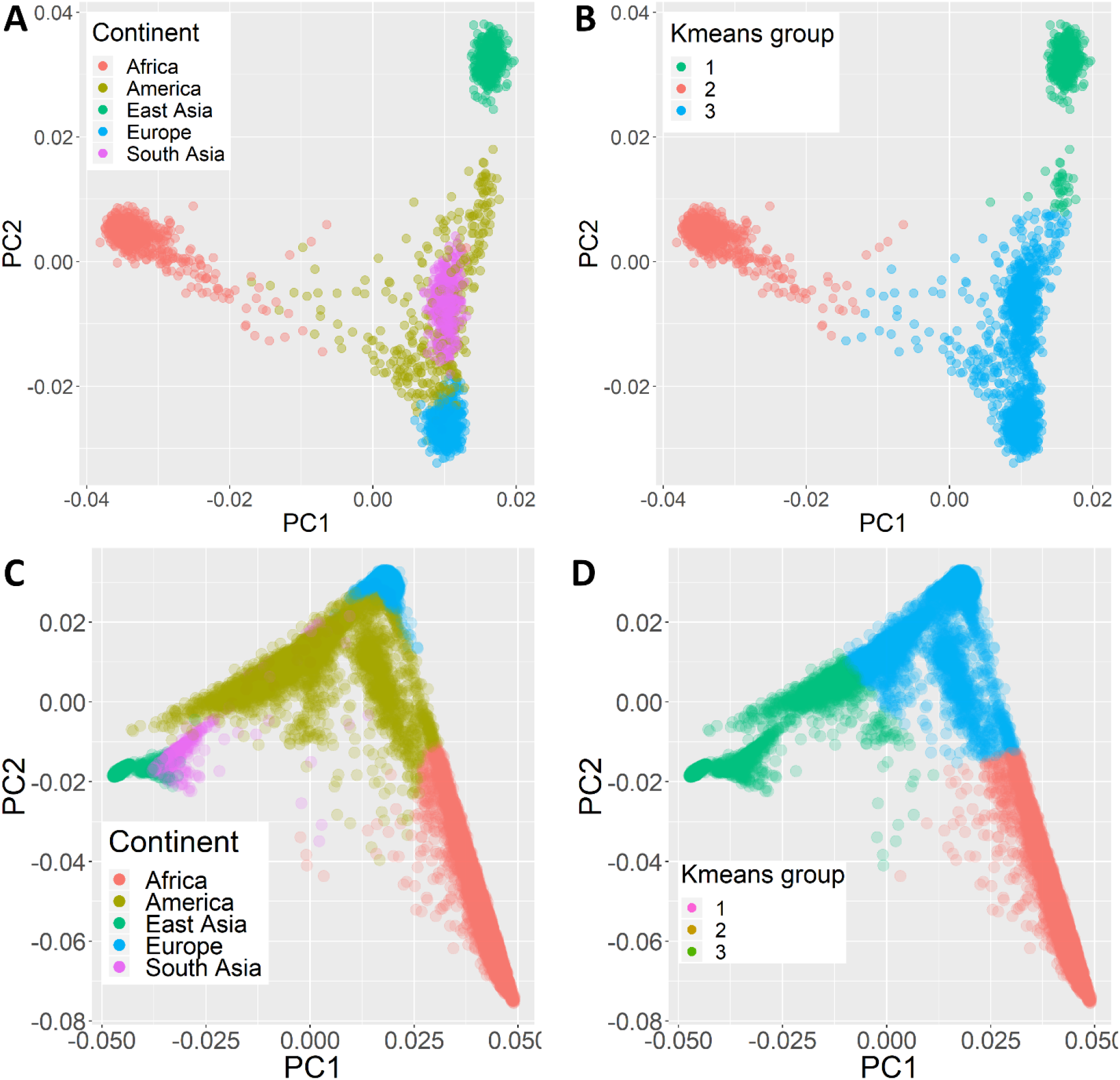
Principal component plots and group assignments for 1000 Genomes and TOPMed samples. Ancestry group assignments for samples in 1000G (A, B) and TOPMed (C, D) samples used either a high-quality ancestry estimation method (A, C) or a crude k-means based method (B, D). In meta-analysis, samples within a group were first analyzed together using the unadjusted test. Then, the group-level results were combined using Stouffer’s method. Meta-analyses using the cruder k-means groupings performed much worse than those using the high-quality ancestry estimates due to population stratification within the cruder groups.

**Figure S10.**
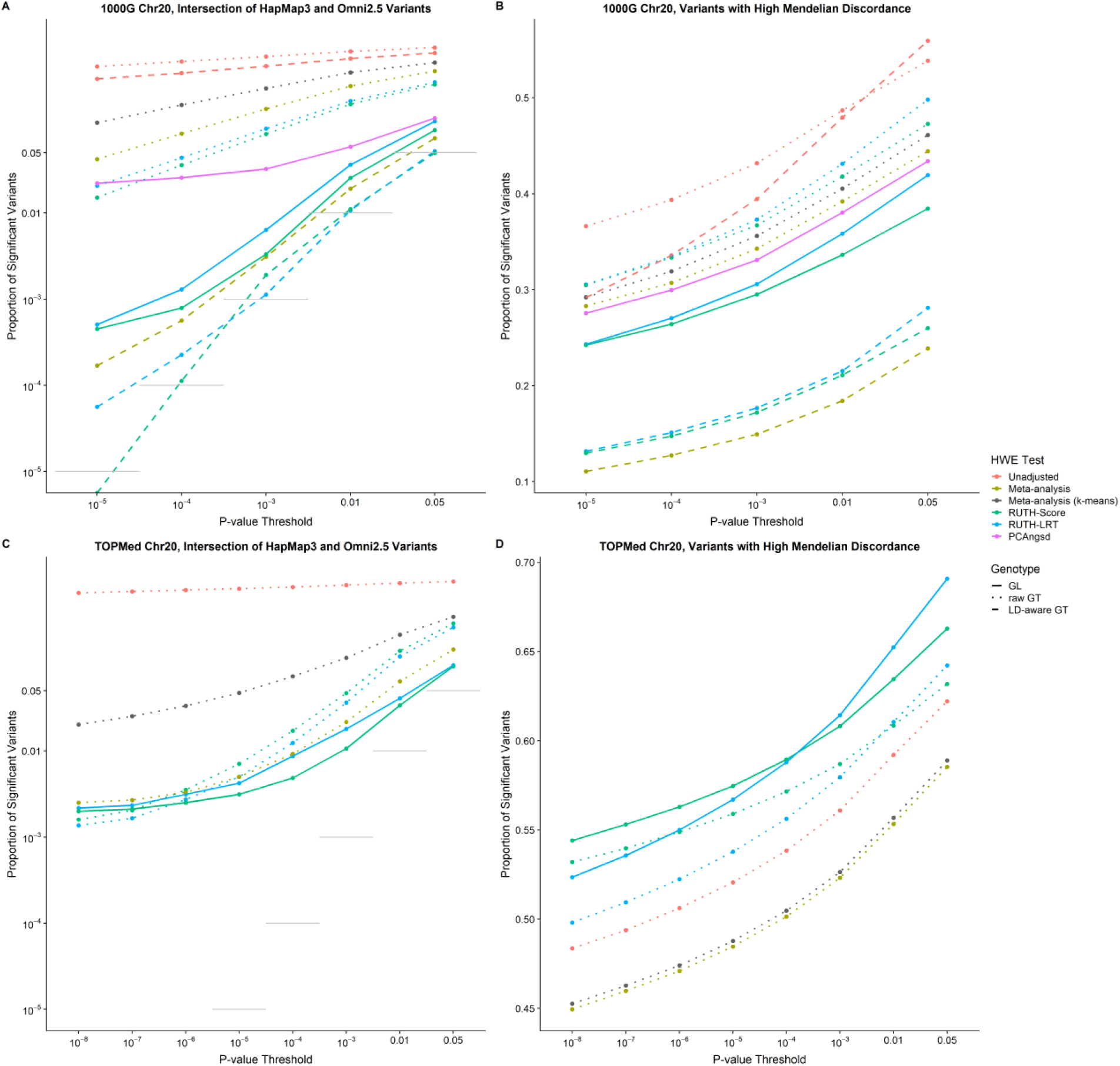
Results of testing 1000G and TOPMed variants with meta-analysis using K-means to generate ancestry groups. We generated three subpopulations for 1000G and TOPMed separately by applying k-means to the first two principal components of each group. Next, we calculated subpopulation-specific HWE statistics, which were converted to Z-scores and combined using Stouffer’s method, using each subpopulation’s size as the weights. (A) K-means-based meta-analysis had much higher false positive rates in 1000G compared to meta-analysis that used more accurate population labels, which (B) confounds its seemingly higher power to discover true positives. (C) We see the same increased false positive rate in K-means-based meta-analysis in TOPMed, but surprisingly (D) it also reduced the power to discover true positives in TOPMed. High-quality ancestry groups can substantially improve the performance of ancestry-based meta-analysis.

**Table S1**

Simulation results for the unadjusted test, meta-analysis, RUTH, and PCAngsd for HWE.

This table can be found at the following link: https://docs.google.com/spreadsheets/d/1zdn7jOWgOMG_wwqwgDD4b1i0a2clGlyNFKmI5xR_DoE/edit?usp=sharing

Results from various HWE tests for simulations with 50,000 variants for 5,000 samples. Samples were generated using a population fixation index (F_ST_) between .01 and .1. “GL” indicates a method using genotype likelihoods, while “GT” indicates a method using best-guess genotypes. Theta denotes deviation from HWE: Theta = 0 indicates no deviation from HWE, Theta < 0 indicates excess heterozygosity, and Theta > 0 indicates heterozygote depletion. When the samples were generated from a single ancestry, meta-analysis and the unadjusted test were identical. *Combined F_ST_ indicates the combined results for F_ST_=.01, .02, .03, .05, and .1. This is available only when the number of ancestries is 1, because F_ST_ should not affect the results with single ancestry, so the results may be combined.

**Table S2.**
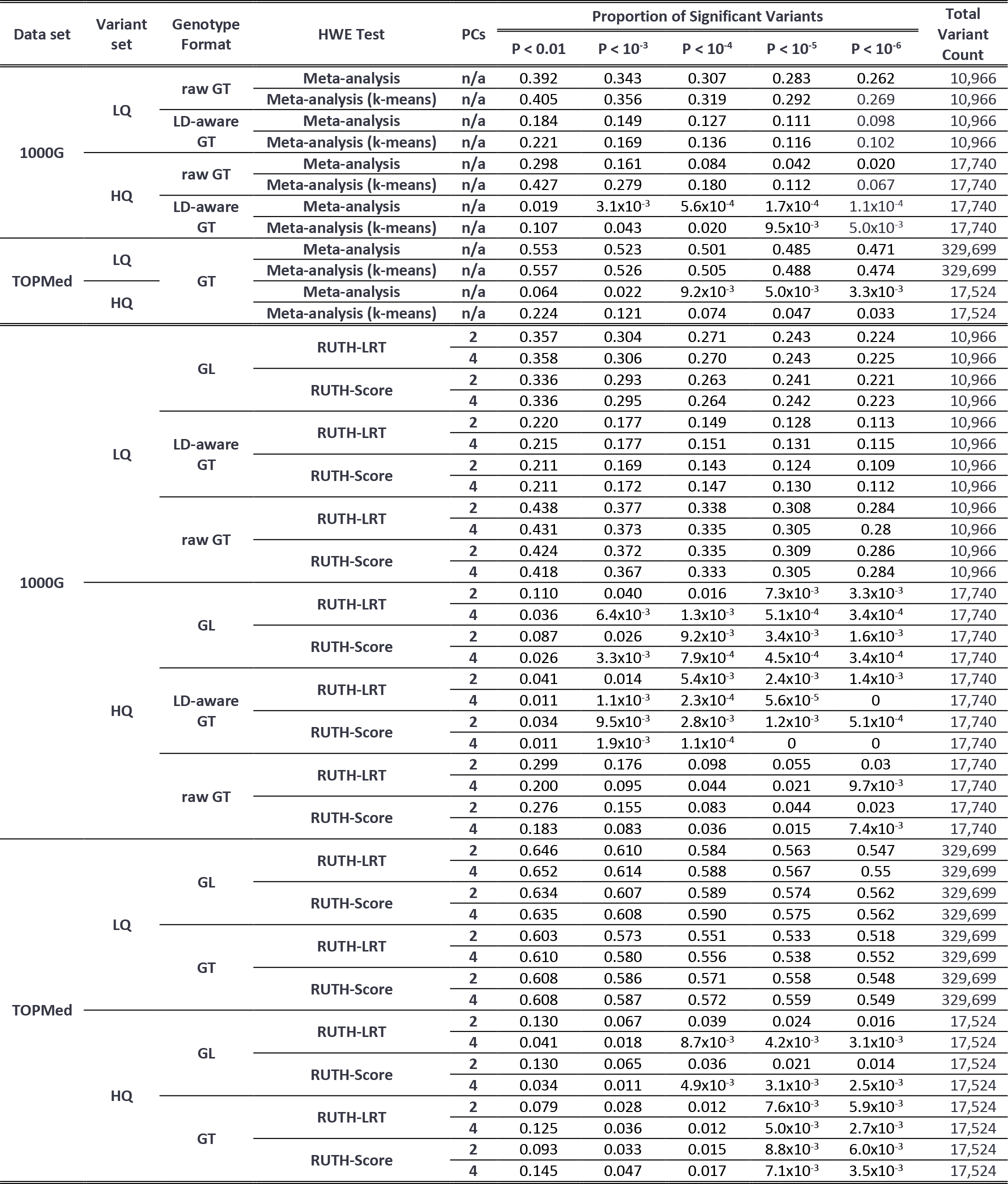
Results from using lower quality ancestry estimations on meta-analysis and RUTH. In both 1000G and TOPMed, the false positive rate was much higher when k-means-based groupings were used for meta-analysis, compared to when high quality ancestry groupings were used. Similarly, the false positive rate was much higher when only 2 PCs were used, compared to when 4 PCs were used. Surprisingly, in TOPMed, using 4 PCs led to both a lower false positive rate and higher true positive rate when compared to using 2 PCs.

**Table S3.** Performance of the unadjusted test, meta-analysis, and RUTH on the subset of TOPMed freeze 5 chromosome 20 variants that are also found in 1000G. For HQ variants, GL-based HWE tests had much better control of false positives than GT-based tests. Conversely, for LQ variants, GT-based HWE tests had a slightly better true positive rate than GL-based tests. Overall, GL-based tests had the best performance when considering the tradeoff between false positives and true positives (Figure S5–6).

| Variant set | Genotype Format | HWE Test | Proportion of Significant Variants |  |  |  |  | Total Variant Count |
| --- | --- | --- | --- | --- | --- | --- | --- | --- |
| | | | $P < 10^{-2}$ | $P < 10^{-3}$ | $P < 10^{-4}$ | $P < 10^{-5}$ | $P < 10^{-6}$ | |
| HQ Variants | raw GT | Unadjusted | 0.890 | 0.842 | 0.800 | 0.766 | 0.736 | 16,924 |
| | raw GT | Meta-analysis | 0.062 | 0.020 | $8.0 \times 10^{-3}$ | $3.8 \times 10^{-3}$ | $2.3 \times 10^{-3}$ | 16,924 |
| | raw GT | RUTH-Score | 0.145 | 0.046 | 0.016 | $6.3 \times 10^{-3}$ | $2.8 \times 10^{-3}$ | 16,924 |
| | GL | RUTH-Score | 0.032 | $9.3 \times 10^{-3}$ | $3.7 \times 10^{-3}$ | $2.0 \times 10^{-3}$ | $1.5 \times 10^{-3}$ | 16,924 |
| | raw GT | RUTH-LRT | 0.125 | 0.035 | 0.011 | $4.2 \times 10^{-3}$ | $1.9 \times 10^{-3}$ | 16,924 |
| | GL | RUTH-LRT | 0.039 | 0.016 | $7.4 \times 10^{-3}$ | $3.1 \times 10^{-3}$ | $2.2 \times 10^{-3}$ | 16,924 |
| LQ Variants | raw GT | Unadjusted | 0.762 | 0.728 | 0.702 | 0.683 | 0.667 | 10,513 |
|  | raw GT | Meta-analysis | 0.649 | 0.616 | 0.592 | 0.575 | 0.560 | 10,513 |
|  | raw GT | RUTH-Score | 0.727 | 0.693 | 0.673 | 0.656 | 0.640 | 10,513 |
|  | GL | RUTH-Score | 0.698 | 0.669 | 0.648 | 0.631 | 0.618 | 10,513 |
|  | raw GT | RUTH-LRT | 0.719 | 0.686 | 0.663 | 0.643 | 0.627 | 10,513 |
|  | GL | RUTH-LRT | 0.693 | 0.662 | 0.639 | 0.621 | 0.605 | 10,513 |

**Table S4**

Simulation results for RUTH tests using 2 vs 4 principal components.

This table can be found at the following link: https://docs.google.com/spreadsheets/d/1Ac9rveZax5Y8NlKQ47wBaJNELqeJkFuNUpa1sNgnsno/edit?usp=sharing We tested the effect of using different numbers of PCs in RUTH on Type I Error (θ = 0) and power (θ ≠ 0) for simulated samples with different numbers of ancestries, fixation indices, sequencing depths, and genotype representations. We simulated 50,000 variants for each combination of simulation parameters.

**Table S5.**
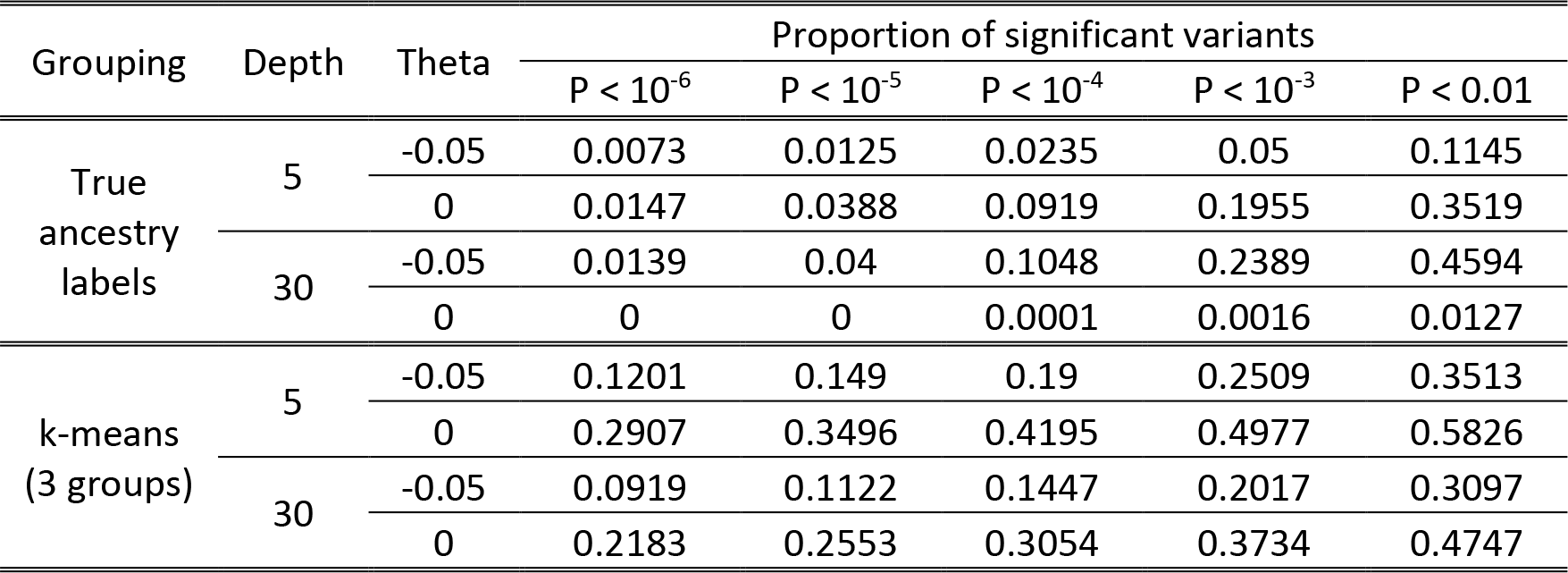
The effect of high vs. low quality subpopulation classification on meta-analysis in simulated samples. We simulated 50,000 variants in 5,000 samples arising from 5 distinct subpopulations (1,000 samples each), at low (5x) and high (30x) depth, with no deviation from HWE (θ = 0) and moderate excess heterozygosity (θ = −0.05). We used one of two different groupings for our samples: for high-quality labels, we used the original true ancestry labels from which we simulated our data; for low-quality labels, we ran k-means classification on the first 2 principal components of genetic variation for all our samples to generate 3 groups. We meta-analyzed all data sets using Stouffer’s method. Type I error rates for low-depth samples were greatly inflated. For high-depth samples, when we used the true ancestry labels, Type I errors were well-controlled, with reasonable power to discover deviations from HWE, while when we used the crude k-means labels, Type I errors were greatly inflated, with surprisingly less power to discover deviations from HWE at less stringent P-value thresholds. These results highlight the importance of high-quality subpopulation classification for meta-analysis.

| Grouping | Depth | Theta | Proportion of significant variants |  |  |  |  |
| --- | --- | --- | --- | --- | --- | --- | --- |
| | | | $P < 10^{-6}$ | $P < 10^{-5}$ | $P < 10^{-4}$ | $P < 10^{-3}$ | $P < 0.01$ |
| True ancestry labels | 5 | -0.05 | 0.0073 | 0.0125 | 0.0235 | 0.05 | 0.1145 |
|  |  | 0 | 0.0147 | 0.0388 | 0.0919 | 0.1955 | 0.3519 |
|  | 30 | -0.05 | 0.0139 | 0.04 | 0.1048 | 0.2389 | 0.4594 |
|  |  | 0 | 0 | 0 | 0.0001 | 0.0016 | 0.0127 |
| k-means (3 groups) | 5 | -0.05 | 0.1201 | 0.149 | 0.19 | 0.2509 | 0.3513 |
|  |  | 0 | 0.2907 | 0.3496 | 0.4195 | 0.4977 | 0.5826 |
|  | 30 | -0.05 | 0.0919 | 0.1122 | 0.1447 | 0.2017 | 0.3097 |
|  |  | 0 | 0.2183 | 0.2553 | 0.3054 | 0.3734 | 0.4747 |

**Table S6.** Comparison of runtimes and memory requirements for RUTH and PCAngsd in simulated and 1000G data. Simulation runtimes for PLINK and RUTH are averaged over 360 runs, across combinations of different simulation parameters. Simulation results for PCAngsd are averaged over 66 runs each for 5x and 30x coverage data. The higher uncertainty in low depth simulated data appears to have led to slower convergence in PCAngsd. All results for 1000G were from single runs. The listed TOPMed runtimes and memory requirements are for single-threaded analyses of all variants.

| Data set | Genotype Format | Software | Test | N | Total Variant Count | Runtime (s) | Memory requirement (MB) |
| --- | --- | --- | --- | --- | --- | --- | --- |
| Simulated | GT | PLINK | Unadjusted | 5,000 | 50,000 | 22 | 10 |
|  | GT | RUTH | RUTH LRT | 5,000 | 50,000 | 348 | 15 |
|  | GL | RUTH | RUTH LRT | 5,000 | 50,000 | 341 | 15 |
|  | GT | RUTH | RUTH Score | 5,000 | 50,000 | 460 | 15 |
|  | GL | RUTH | RUTH Score | 5,000 | 50,000 | 469 | 15 |
| Simulated (5x) | GL | PCAngsd | PCAngsd | 5,000 | 50,000 | 6,068 | 6,946 |
| Simulated (30x) | GL | PCAngsd | PCAngsd | 5,000 | 50,000 | 5,337 | 6,872 |
| 1000G | GT | PLINK | Unadjusted | 2,504 | 28,706 | 2 | 8 |
|  | GL | RUTH | RUTH LRT | 2,504 | 28,706 | 147 | 14 |
|  | GT | RUTH | RUTH LRT | 2,504 | 28,706 | 96 | 13 |
|  | GL | RUTH | RUTH Score | 2,504 | 28,706 | 216 | 14 |
|  | GT | RUTH | RUTH Score | 2,504 | 28,706 | 177 | 13 |
|  | GL | PCAngsd | PCAngsd | 2,504 | 28,660 | 4,105 | 2,073 |
| TOPMed | GT | RUTH | RUTH LRT | 53,831 | 347,223 | 158,731 | 57 |
|  | GL | RUTH | RUTH LRT | 53,831 | 347,223 | 196,169 | 57 |

**Table S7.** Sample contributions from each of the participating TOPMed studies.

| TOPMed Study Name | TOPMed Accession | Sample Size |
| --- | --- | --- |
| Genetics of Cardiometabolic Health in the Amish | phs000956 | 1,025 |
| Trans-Omics for Precision Medicine Whole Genome Sequencing Project: ARIC | phs001211 | 3,585 |
| The Genetics and Epidemiology of Asthma in Barbados | phs001143 | 944 |
| Cleveland Clinic Atrial Fibrillation Study | phs001189 | 328 |
| The Cleveland Family Study (WGS) | phs000954 | 919 |
| Cardiovascular Health Study | phs001368 | 69 |
| Genetic Epidemiology of COPD (COPDGene) in the TOPMed Program | phs000951 | 8,733 |
| The Genetic Epidemiology of Asthma in Costa Rica | phs000988 | 1,040 |
| Diabetes Heart Study African American Coronary Artery Calcification (AA CAC) | phs001412 | 322 |
| Whole Genome Sequencing and Related Phenotypes in the Framingham Heart Study | phs000974 | 3,725 |
| Genes-environments and Admixture in Latino Asthmatics (GALA II) Study | phs000920 | 912 |
| GeneSTAR (Genetic Study of Atherosclerosis Risk) | phs001218 | 1,633 |
| Genetic Epidemiology Network of Arteriopathy (GENOA) | phs001345 | 1,069 |
| Genetic Epidemiology Network of Salt Sensitivity (GenSalt) | phs001217 | 1,680 |
| Genetics of Lipid Lowering Drugs and Diet Network (GOLDN) | phs001359 | 892 |
| Heart and Vascular Health Study (HVH) | phs000993 | 64 |
| HyperGEN - Genetics of Left Ventricular (LV) Hypertrophy | phs001293 | 1,752 |
| Jackson Heart Study | phs000964 | 3,074 |
| Whole Genome Sequencing of Venous Thromboembolism (WGS of VTE) | phs001402 | 1,250 |
| MESA and MESA Family AA-CAC | phs001416 | 4,804 |
| MGH Atrial Fibrillation Study | phs001062 | 916 |
| Partners HealthCare Biobank | phs001024 | 109 |
| San Antonio Family Heart Study (WGS) | phs001215 | 1,478 |
| Study of African Americans, Asthma, Genes and Environment (SAGE) Study | phs000921 | 450 |
| African American Sarcoidosis Genetics Resource | phs001207 | 606 |
| Genome-wide Association Study of Adiposity in Samoans | phs000972 | 1,198 |
| The Vanderbilt AF Ablation Registry | phs000997 | 154 |
| The Vanderbilt Atrial Fibrillation Registry | phs001032 | 1016 |
| Novel Risk Factors for the Development of Atrial Fibrillation in Women | phs001040 | 97 |
| Women's Health Initiative (WHI) | phs001237 | 9,984 |
| Total |  | 53,831 |

**Table S8.**
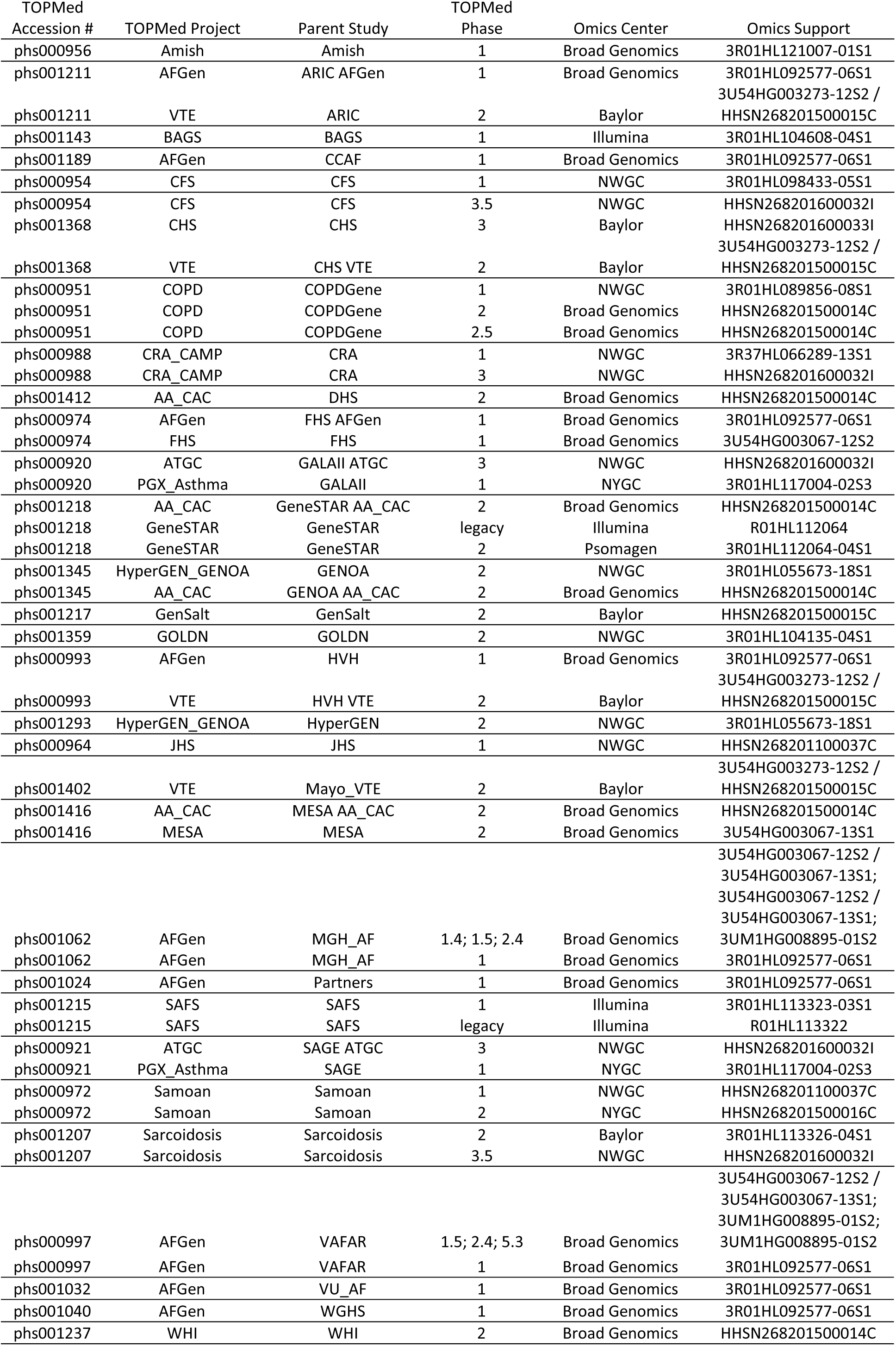
TOPMed acknowledgements for omics support.

### File S1

#### TOPMed Study Acknowledgements

##### NHLBI TOPMed: Genetics of Cardiometabolic Health in the Amish

The Amish studies upon which these data are based were supported by NIH grants R01 AG18728, U01 HL072515, R01 HL088119, R01 HL121007, and P30 DK072488. See publication: PMID: 18440328

##### NHLBI TOPMed: Trans-Omics for Precision Medicine Whole Genome Sequencing Project: ARIC

Genome Sequencing for “NHLBI TOPMed: Atherosclerosis Risk in Communities (ARIC)” (phs001211) was performed at the Baylor College of Medicine Human Genome Sequencing Center (HHSN268201500015C and 3U54HG003273-12S2) and the Broad Institute of MIT and Harvard (3R01HL092577-06S1).

The Atherosclerosis Risk in Communities study has been funded in whole or in part with Federal funds from the National Heart, Lung, and Blood Institute, National Institutes of Health, Department of Health and Human Services (contract numbers HHSN268201700001I, HHSN268201700002I, HHSN268201700003I, HHSN268201700004I and HHSN268201700005I).

The authors thank the staff and participants of the ARIC study for their important contributions.

##### NHLBI TOPMed: The Genetics and Epidemiology of Asthma in Barbados

The Genetics and Epidemiology of Asthma in Barbados is supported by National Institutes of Health (NIH) National Heart, Lung, Blood Institute TOPMed (R01 HL104608-S1) and: R01 AI20059, K23 HL076322, and RC2 HL101651. For the specific cohort descriptions and descriptions regarding the collection of phenotype data can be found at: https://www.nhlbiwgs.org/group/bags-asthma. The authors wish to give special recognition to the individual study participants who provided biological samples and or data, without their support in research none of this would be possible.

##### NHLBI TOPMed: Cleveland Clinic Atrial Fibrillation Study

The research reported in this article was supported by grants from the National Institutes of Health (NIH) National Heart, Lung, and Blood Institute grants R01 HL090620 and R01 HL111314, the NIH National Center for Research Resources for Case Western Reserve University and the Cleveland Clinic Clinical and Translational Science Award (CTSA) UL1-RR024989, the Department of Cardiovascular Medicine philanthropic research fund, Heart and Vascular Institute, Cleveland Clinic, the Fondation Leducq grant 07-CVD 03, and The Atrial Fibrillation Innovation Center, State of Ohio.

##### NHLBI TOPMed: The Cleveland Family Study (WGS)

Support for the Cleveland Family Study was provided by NHLBI grant numbers R01 HL46380, R01 HL113338 and R35 HL135818.

##### NHLBI TOPMed: Cardiovascular Health Study

This research was supported by contracts HHSN268201200036C, HHSN268200800007C, HHSN268201800001C, N01-HC85079, N01-HC-85080, N01-HC-85081, N01-HC-85082, N01-HC-85083, N01-HC-85084, N01-HC-85085, N01-HC-85086, N01-HC-35129, N01-HC-15103, N01-HC-55222, N01-HC-75150, N01-HC-45133, and N01-HC-85239; grant numbers U01 HL080295, U01 HL130114 and R01 HL059367 from the National Heart, Lung, and Blood Institute, and R01 AG023629 from the National Institute on Aging, with additional contributions from the National Institute of Neurological Disorders and Stroke. A full list of principal CHS investigators and institutions can be found at https://chs-nhlbi.org/pi. Its content is solely the responsibility of the authors and does not necessarily represent the official views of the National Institutes of Health.

##### NHLBI TOPMed: Genetic Epidemiology of COPD (COPDGene) in the TOPMed Program

This research used data generated by the COPDGene study, which was supported by NIH Award Number U01 HL089897 and Award Number U01 HL089856 from the National Heart, Lung, and Blood Institute. The content is solely the responsibility of the authors and does not necessarily represent the official views of the National Heart, Lung, and Blood Institute or the National Institutes of Health.

The COPDGene project is also supported by the COPD Foundation through contributions made to an Industry Advisory Board comprised of AstraZeneca, Boehringer Ingelheim, GlaxoSmithKline, Novartis, Pfizer, Siemens and Sunovion.

##### NHLBI TOPMed: The Genetic Epidemiology of Asthma in Costa Rica

This study was supported by NHLBI grants R37 HL066289 and P01 HL132825. We wish to acknowledge the investigators at the Channing Division of Network Medicine at Brigham and Women’s Hospital, the investigators at the Hospital Nacional de Niños in San José, Costa Rica and the study subjects and their extended family members who contributed samples and genotypes to the study, and the NIH/NHLBI for its support in making this project possible.

##### NHLBI TOPMed: Diabetes Heart Study African American Coronary Artery Calcification (AA CAC)

This work was supported by R01 HL92301, R01 HL67348, R01 NS058700, R01 AR48797, R01 DK071891, R01 AG058921, the General Clinical Research Center of the Wake Forest University School of Medicine (M01 RR07122, F32 HL085989), the American Diabetes Association, and a pilot grant from the Claude Pepper Older Americans Independence Center of Wake Forest University Health Sciences (P60 AG10484).

##### NHLBI TOPMed: Whole Genome Sequencing and Related Phenotypes in the Framingham Heart Study

The Framingham Heart Study (FHS) is a prospective cohort study of 3 generations of subjects who have been followed up to 65 years to evaluate risk factors for cardiovascular disease.13-16 Its large sample of ~15,000 men and women who have been extensively phenotyped with repeated examinations make it ideal for the study of genetic associations with cardiovascular disease risk factors and outcomes. DNA samples have been collected and immortalized since the mid-1990s and are available on ~8000 study participants in 1037 families. These samples have been used for collection of GWAS array data and exome chip data in nearly all with DNA samples, and for targeted sequencing, deep exome sequencing and light coverage whole genome sequencing in limited numbers. Additionally, mRNA and miRNA expression data, DNA methylation data, metabolomics and other ‘omics data are available on a sizable portion of study participants. This project will focus on deep whole genome sequencing (mean 30X coverage) in ~4100 subjects and imputed to all with GWAS array data to more fully understand the genetic contributions to cardiovascular, lung, blood and sleep disorders.

The FHS acknowledges the support of contracts NO1-HC-25195 and HHSN268201500001I from the National Heart, Lung, and Blood Institute and grant supplement R01 HL092577-06S1 for this research. We also acknowledge the dedication of the FHS study participants without whom this research would not be possible.

##### NHLBI TOPMed: Genes-environments and Admixture in Latino Asthmatics (GALA II) Study

The Genes-environments and Admixture in Latino Americans (GALA II) Study was supported by the National Heart, Lung, and Blood Institute of the National Institute of Health (NIH) grants R01HL117004 and X01HL134589; study enrollment supported by the Sandler Family Foundation, the American Asthma Foundation, the RWJF Amos Medical Faculty Development Program, Harry Wm. and Diana V. Hind Distinguished Professor in Pharmaceutical Sciences II and the National Institute of Environmental Health Sciences grant R01ES015794.

The GALA II study collaborators include Shannon Thyne, UCSF; Harold J. Farber, Texas Children’s Hospital; Denise Serebrisky, Jacobi Medical Center; Rajesh Kumar, Lurie Children’s Hospital of Chicago; Emerita Brigino-Buenaventura, Kaiser Permanente; Michael A. LeNoir, Bay Area Pediatrics; Kelley Meade, UCSF Benioff Children’s Hospital, Oakland; William Rodriguez-Cintron, VA Hospital, Puerto Rico; Pedro C. Avila, Northwestern University; Jose R. Rodriguez-Santana, Centro de Neumologia Pediatrica; Luisa N. Borrell, City University of New York; Adam Davis, UCSF Benioff Children’s Hospital, Oakland; Saunak Sen, University of Tennessee and Fred Lurmann, Sonoma Technologies, Inc.

The authors acknowledge the families and patients for their participation and thank the numerous health care providers and community clinics for their support and participation in GALA II. In particular, the authors thank study coordinator Sandra Salazar; the recruiters who obtained the data: Duanny Alva, MD, Gaby Ayala-Rodriguez, Lisa Caine, Elizabeth Castellanos, Jaime Colon, Denise DeJesus, Blanca Lopez, Brenda Lopez, MD, Louis Martos, Vivian Medina, Juana Olivo, Mario Peralta, Esther Pomares, MD, Jihan Quraishi, Johanna Rodriguez, Shahdad Saeedi, Dean Soto, Ana Taveras; and the lab researcher Celeste Eng who processed the biospecimens.

##### NHLBI TOPMed: Genetic Epidemiology Network of Arteriopathy (GENOA)

Support for GENOA was provided by the National Heart, Lung and Blood Institute (HL054457, HL054464, HL054481, HL119443, and HL087660) of the National Institutes of Health. WGS for “NHLBI TOPMed: Genetic Epidemiology Network of Arteriopathy” (phs001345) was performed at the Mayo Clinic Genotyping Core, the DNA Sequencing and Gene Analysis Center at the University of Washington (3R01HL055673-18S1), and the Broad Institute (HHSN268201500014C) for their genotyping and sequencing services. We would like to thank the GENOA participants.

##### NHLBI TOPMed: Genetic Epidemiology Network of Salt Sensitivity (GenSalt)

The Genetic Epidemiology Network of Salt-Sensitivity (GenSalt) was supported by research grants (U01HL072507, R01HL087263, and R01HL090682) from the National Heart, Lung, and Blood Institute, National Institutes of Health, Bethesda, MD.

##### NHLBI TOPMed: Genetics of Lipid Lowering Drugs and Diet Network (GOLDN)

GOLDN biospecimens, baseline phenotype data, and intervention phenotype data were collected with funding from the National Heart, Lung and Blood Institute (NHLBI) grant U01 HL072524. Whole-genome sequencing in GOLDN was funded by NHLBI grant R01 HL104135 and supplement R01 HL104135-04S1.

##### NHLBI TOPMed: Heart and Vascular Health Study (HVH)

The research reported in this article was supported by grants HL068986, HL085251, HL095080, and HL073410 from the National Heart, Lung, and Blood Institute.

##### NHLBI TOPMed: Hypertension Genetic Epidemiology Network (HyperGEN)

The HyperGEN Study is part of the National Heart, Lung, and Blood Institute (NHLBI) Family Blood Pressure Program; collection of the data represented here was supported by grants U01 HL054472 (MN Lab), U01 HL054473 (DCC), U01 HL054495 (AL FC), and U01 HL054509 (NC FC). The HyperGEN: Genetics of Left Ventricular Hypertrophy Study was supported by NHLBI grant R01 HL055673 with whole-genome sequencing made possible by supplement −18S1.

##### NHLBI TOPMed: The Jackson Heart Study

The Jackson Heart Study (JHS) is supported and conducted in collaboration with Jackson State University (HHSN268201800013I), Tougaloo College (HHSN268201800014I), the Mississippi State Department of Health (HHSN268201800015I/HHSN26800001) and the University of Mississippi Medical Center (HHSN268201800010I, HHSN268201800011I and HHSN268201800012I) contracts from the National Heart, Lung, and Blood Institute (NHLBI) and the National Institute for Minority Health and Health Disparities (NIMHD). The authors also wish to thank the staffs and participants of the JHS.

##### NHLBI TOPMed: Multi-Ethnic Study of Atherosclerosis

MESA and the MESA SHARe projects are conducted and supported by the National Heart, Lung, and Blood Institute (NHLBI) in collaboration with MESA investigators. Support for MESA is provided by contracts 75N92020D00001, HHSN268201500003I, N01-HC-95159, 75N92020D00005, N01-HC-95160, 75N92020D00002, N01-HC-95161, 75N92020D00003, N01-HC-95162, 75N92020D00006, N01-HC-95163, 75N92020D00004, N01-HC-95164, 75N92020D00007, N01-HC-95165, N01-HC-95166, N01-HC-95167, N01-HC-95168, N01-HC-95169, UL1-TR-000040, UL1-TR-001079, and UL1-TR-001420. Also supported by the National Center for Advancing Translational Sciences, CTSI grant UL1TR001881, and the National Institute of Diabetes and Digestive and Kidney Disease Diabetes Research Center (DRC) grant DK063491 to the Southern California Diabetes Endocrinology Research Center.

##### NHLBI TOPMed: Whole Genome Sequencing of Venous Thromboembolism (WGS of VTE)

Funded in part by grants from the National Institutes of Health, National Heart, Lung, and Blood Institute (HL66216 and HL83141) and the National Human Genome Research Institute (HG04735).

##### NHLBI TOPMed: MGH Atrial Fibrillation Study

This work was supported by the Fondation Leducq (14CVD01), and by grants from the National Institutes of Health to Dr. Ellinor (1RO1HL092577, R01HL128914, K24HL105780). This work was also supported by a grant from the American Heart Association to Dr. Ellinor (18SFRN34110082). Dr. Lubitz is supported by NIH grant 1R01HL139731 and AHA 18SFRN34250007.

##### NHLBI TOPMed: Partners HealthCare Biobank

We thank the Broad Institute for generating high-quality sequence data supported by the NHLBI grant 3R01HL092577-06S1 to Dr. Patrick Ellinor. The datasets used in this manuscript were obtained from dbGaP at http://www.ncbi.nlm.nih.gov/gap through dbGaP accession number phs001024.

##### NHLBI TOPMed: Study of African Americans, Asthma, Genes and Environment (SAGE)

The Study of African Americans, Asthma, Genes and Environments (SAGE) was supported by the National Heart, Lung, and Blood Institute of the National Institute of Health (NIH) grants R01HL117004 and X01HL134589; study enrollment supported by the Sandler Family Foundation, the American Asthma Foundation, the RWJF Amos Medical Faculty Development Program, Harry Wm. and Diana V. Hind Distinguished Professor in Pharmaceutical Sciences II. The SAGE study collaborators include Harold J. Farber, Texas Children’s Hospital; Emerita Brigino-Buenaventura, Kaiser Permanente; Michael A. LeNoir, Bay Area Pediatrics; Kelley Meade, UCSF Benioff Children’s Hospital, Oakland; Luisa N. Borrell, City University of New York; Adam Davis, UCSF Benioff Children’s Hospital, Oakland and Fred Lurmann, Sonoma Technologies, Inc.

The authors acknowledge the families and patients for their participation and thank the numerous health care providers and community clinics for their support and participation in SAGE. In particular, the authors thank study coordinator Sandra Salazar; the recruiters who obtained the data: Lisa Caine, Elizabeth Castellanos, Brenda Lopez, MD, Shahdad Saeedi; and the lab researcher Celeste Eng who processed the biospecimens.

E.G.B was supported by National Heart, Lung, and Blood Institute (NHLBI): U01HL138626, R01HL117004, R01HL128439, R01HL135156, X01HL134589, R01HL141992, R01HL141845; the National Human Genome Research Institute (NHGRI): U01HG009080; the National Institute of Environmental Health Sciences (NIEHS): R01ES015794, R21ES24844; the National Institute on Minority Health and Health Disparities (NIMHD): P60MD006902, R01MD010443, RL5GM118984,R56MD013312; the Eunice Kennedy Shriver National Institute of Child Health and Human Development (NICHD): R01HD085993; and the Tobacco-Related Disease Research Program (TRDRP): 24RT-0025 and 27IR-0030.

##### NHLBI TOPMed: San Antonio Family Heart Study (WGS)

Collection of the San Antonio Family Study data was supported in part by National Institutes of Health (NIH) grants R01 HL045522, MH078143, MH078111 and MH083824; and whole genome sequencing of SAFS subjects was supported by U01 DK085524 and R01 HL113323. We are very grateful to the participants of the San Antonio Family Study for their continued involvement in our research programs.

##### NHLBI TOPMed: The Samoan Obesity, Lifestyle and Genetic Adaptations Study (OLaGA) Group

Financial support for the Samoan Obesity, Lifestyle and Genetic Adaptations Study (OLaGA) Group comes from the U.S. National Institutes of Health Grant R01-HL093093 and R01-HL133040. We acknowledge the assistance of the Samoa Ministry of Health and the Samoa Bureau of Statistics for their guidance and support in the conduct of this study. We thank the local village officials for their help and the participants for their generosity. The following publication describes the origin of the dataset: Hawley NL, Minster RL, Weeks DE, Viali S, Reupena MS, Sun G, Cheng H, Deka R, McGarvey ST. Prevalence of Adiposity and Associated Cardiometabolic Risk Factors in the Samoan Genome-Wide Association Study. Am J Human Biol 2014. 26: 491-501. DOI: 10.1002/jhb.22553. PMID: 24799123.

Our study name: ‘The Samoan Obesity, Lifestyle and Genetic Adaptations Study (OLaGA) Group’.

Ranjan Deka, Department of Environmental and Public Health Sciences, College of Medicine, University of Cincinnati, Cincinnati, OH 45267-0056.

Nicola L Hawley, Department of Epidemiology (Chronic Disease), School of Public Health, Yale University, New Haven, CT 06520-0834.

Stephen T McGarvey, International Health Institute, Department of Epidemiology, School of Public Health, and Department of Anthropology, Brown University. 02912.

Ryan L Minster, Department of Human Genetics and Department of Biostatistics, University of Pittsburgh, Pittsburgh, PA 15261.

Take Naseri, Ministry of Health, Government of Samoa, Apia, Samoa.

Muagututi’a Sefuiva Reupena, Lutia I Puava Ae Mapu I Fagalele, Apia, Samoa.

Daniel E Weeks, Department of Human Genetics and Department of Biostatistics, University of Pittsburgh, Pittsburgh, PA 15261.

##### NHLBI TOPMed: The Vanderbilt AF Ablation Registry

The research reported in this article was supported by grants from the American Heart Association to Dr. Shoemaker (11CRP742009), Dr. Darbar (EIA 0940116N), and grants from the National Institutes of Health (NIH) to Dr. Darbar (R01 HL092217), and Dr. Roden (U19 HL65962, and UL1 RR024975). The project was also supported by a CTSA award (UL1 TR00045) from the National Center for Advancing Translational Sciences. Its contents are solely the responsibility of the authors and do not necessarily represent the official views of the National Center for Advancing Translational Sciences or the NIH.

##### NHLBI TOPMed: The Vanderbilt Atrial Fibrillation Registry

The research reported in this article was supported by grants from the American Heart Association to Dr. Darbar (EIA 0940116N), and grants from the National Institutes of Health (NIH) to Dr. Darbar (HL092217), and Dr. Roden (U19 HL65962, and UL1 RR024975). This project was also supported by CTSA award (UL1TR000445) from the National Center for Advancing Translational Sciences. Its contents are solely the responsibility of the authors and do not necessarily represent the official views of the National Center for Advancing Translational Sciences of the NIH.

##### NHLBI TOPMed: Novel Risk Factors for the Development of Atrial Fibrillation in Women

The Women’s Genome Health Study (WGHS) is supported by HL 043851 and HL099355 from the National Heart, Lung, and Blood Institute and CA 047988 from the National Cancer Institute, the Donald W. Reynolds Foundation with collaborative scientific support and funding for genotyping provided by Amgen. AF endpoint confirmation was supported by HL-093613 and a grant from the Harris Family Foundation and Watkin’s Foundation.

##### NHLBI TOPMed: Women’s Health Initiative (WHI)

The WHI program is funded by the National Heart, Lung, and Blood Institute, National Institutes of Health, U.S. Department of Health and Human Services through contracts HHSN268201600018C, HHSN268201600001C, HHSN268201600002C, HHSN268201600003C, and HHSN268201600004C.

##### NHLBI TOPMed: GeneSTAR (Genetic Study of Atherosclerosis Risk)

The Johns Hopkins Genetic Study of Atherosclerosis Risk (GeneSTAR) was supported by grants from the National Institutes of Health through the National Heart, Lung, and Blood Institute (U01HL72518, HL087698, HL112064) and by a grant from the National Center for Research Resources (M01-RR000052) to the Johns Hopkins General Clinical Research Center. We would like to thank the participants and families of GeneSTAR and our dedicated staff for all their sacrifices.

##### NHLBI TOPMed: Genetics of Sarcoidosis in African Americans (Sarcoidosis)

National Institutes of Health (R01HL113326, P30 GM110766-01)

## Notes

### Competing Interest Statement

The authors have declared no competing interest.

### Summary of Updates

Corrected the order of consortium authorships.

## REFERENCES

Balding, D. J., 2003 Likelihood-based inference for genetic correlation coefficients. Theor Popul Biol 63: 221–230.

Balding, D. J., and R. A. Nichols, 1995 A Method for Quantifying Differentiation between Populations at Multi-Allelic Loci and Its Implications for Investigating Identity and Paternity. Genetica 96: 3–12.

Bycroft, C., C. Freeman, D. Petkova, G. Band, L. T. Elliott et al., 2018 The UK Biobank resource with deep phenotyping and genomic data. Nature 562: 203–209.

Danecek, P., A. Auton, G. Abecasis, C. A. Albers, E. Banks et al., 2011 The variant call format and VCFtools. Bioinformatics 27: 2156–2158.

Delaneau, O., J. F. Zagury and J. Marchini, 2013 Improved whole-chromosome phasing for disease and population genetic studies. Nat Methods 10: 5–6.

Dempster, A. P., N. M. Laird and D. B. Rubin, 1977 Maximum Likelihood from Incomplete Data Via the EM Algorithm. Journal of the Royal Statistical Society: Series B (Methodological) 39: 1–22.

Dryden, I. L., and K. V. Mardia, 1998 Statistical shape analysis. John Wiley & Sons, Chichester; New York.

Ewing, B., and P. Green, 1998 Base-calling of automated sequencer traces using phred. II. Error probabilities. Genome Res 8: 186–194.

Fritsche, L. G., W. Igl, J. N. Bailey, F. Grassmann, S. Sengupta et al., 2016 A large genome-wide association study of age-related macular degeneration highlights contributions of rare and common variants. Nat Genet 48: 134–143.

Hao, W., M. Song and J. D. Storey, 2016 Probabilistic models of genetic variation in structured populations applied to global human studies. Bioinformatics 32: 713–721.

Hao, W., and J. D. Storey, 2019 Extending Tests of Hardy-Weinberg Equilibrium to Structured Populations. Genetics 213: 759–770.

Hardy, G. H., 1908 Mendelian Proportions in a Mixed Population. Science 28: 49–50.

Holsinger, K. E., 1999 Analysis of Genetic Diversity in Geographically Structured Populations: A Bayesian Perspective. Hereditas 130: 245–255.

Holsinger, K. E., P. O. Lewis and D. K. Dey, 2002 A Bayesian approach to inferring population structure from dominant markers. Mol Ecol 11: 1157–1164.

Jin, Y., A. A. Schaffer, M. Feolo, J. B. Holmes and B. L. Kattman, 2019 GRAF-pop: A Fast Distance-Based Method To Infer Subject Ancestry from Multiple Genotype Datasets Without Principal Components Analysis. G3 (Bethesda) 9: 2447–2461.

Jun, G., M. Flickinger, K. N. Hetrick, J. M. Romm, K. F. Doheny et al., 2012 Detecting and estimating contamination of human DNA samples in sequencing and array-based genotype data. Am J Hum Genet 91: 839–848.

Kuhn, R. M., D. Haussler and W. J. Kent, 2013 The UCSC genome browser and associated tools. Brief Bioinform 14: 144–161.

Laurie, C. C., K. F. Doheny, D. B. Mirel, E. W. Pugh, L. J. Bierut et al., 2010 Quality control and quality assurance in genotypic data for genome-wide association studies. Genet Epidemiol 34: 591–602.

Li, J. Z., D. M. Absher, H. Tang, A. M. Southwick, A. M. Casto et al., 2008 Worldwide human relationships inferred from genome-wide patterns of variation. Science 319: 1100–1104.

Li, M., and C. Li, 2008 Assessing departure from Hardy-Weinberg equilibrium in the presence of disease association. Genet Epidemiol 32: 589–599.

Locke, A. E., B. Kahali, S. I. Berndt, A. E. Justice, T. H. Pers et al., 2015 Genetic studies of body mass index yield new insights for obesity biology. Nature 518: 197–206.

McCarroll, S. A., T. N. Hadnott, G. H. Perry, P. C. Sabeti, M. C. Zody et al., 2006 Common deletion polymorphisms in the human genome. Nat Genet 38: 86–92.

Meisner, J., and A. Albrechtsen, 2019 Testing for Hardy-Weinberg Equilibrium in Structured Populations using Genotype or Low-Depth NGS Data. Mol Ecol Resour.

Mosteller, F., and R. A. Fisher, 1948 Questions and Answers. The American Statistician 2: 30–31.

Nielsen, D. M., M. G. Ehm and B. S. Weir, 1998 Detecting marker-disease association by testing for Hardy-Weinberg disequilibrium at a marker locus. Am J Hum Genet 63: 1531–1540.

Nielsen, R., J. S. Paul, A. Albrechtsen and Y. S. Song, 2011 Genotype and SNP calling from nextgeneration sequencing data. Nat Rev Genet 12: 443–451.

Price, A. L., N. J. Patterson, R. M. Plenge, M. E. Weinblatt, N. A. Shadick et al., 2006 Principal components analysis corrects for stratification in genome-wide association studies. Nat Genet 38: 904–909.

Purcell, S., B. Neale, K. Todd-Brown, L. Thomas, M. A. Ferreira et al., 2007 PLINK: a tool set for whole-genome association and population-based linkage analyses. Am J Hum Genet 81: 559–575.

Rohlfs, R. V., and B. S. Weir, 2008 Distributions of Hardy-Weinberg equilibrium test statistics. Genetics 180: 1609–1616.

Rosenberg, N. A., J. K. Pritchard, J. L. Weber, H. M. Cann, K. K. Kidd et al., 2002 Genetic structure of human populations. Science 298: 2381–2385.

Sha, Q., and S. Zhang, 2011 A test of Hardy-Weinberg equilibrium in structured populations. Genet Epidemiol 35: 671–678.

Stouffer, S. A., 1949 The American soldier. Princeton University Press, Princeton,.

Stouffer, S. A., E. A. Suchman, L. C. DeVinney, S. A. Star and R. M. Williams Jr, 1949 The American soldier: Adjustment during army life. (Studies in social psychology in World War II), Vol. 1.

Taliun, D., D. N. Harris, M. D. Kessler, J. Carlson, Z. A. Szpiech et al., 2019 Sequencing of 53,831 diverse genomes from the NHLBI TOPMed Program. bioRxiv.

The 1000 Genomes Project Consortium, A. Auton, L. D. Brooks, R. M. Durbin, E. P. Garrison et al., 2015 A global reference for human genetic variation. Nature 526: 68–74.

The International HapMap Consortium, D. M. Altshuler, R. A. Gibbs, L. Peltonen, D. M. Altshuler et al., 2010 Integrating common and rare genetic variation in diverse human populations. Nature 467: 52–58.

Van Oosterhout, C., W. F. Hutchinson, D. P. M. Wills and P. Shipley, 2004 MICRO-CHECKER: software for identifying and correcting genotyping errors in microsatellite data. Molecular Ecology Notes 4: 535–538.

Wang, C., Z. A. Szpiech, J. H. Degnan, M. Jakobsson, T. J. Pemberton et al., 2010 Comparing spatial maps of human population-genetic variation using Procrustes analysis. Stat Appl Genet Mol Biol 9: Article 13.

Waples, R. S., 2015 Testing for Hardy-Weinberg proportions: have we lost the plot? J Hered 106: 1–19.

Weinberg, W., 1908 Uber den nachweis der vererbung beim menschen. Jh. Ver. vaterl. Naturk. Wurttemb. 64: 369–382.

Wigginton, J. E., D. J. Cutler and G. R. Abecasis, 2005 A note on exact tests of Hardy-Weinberg equilibrium. Am J Hum Genet 76: 887–893.

Yang, W. Y., J. Novembre, E. Eskin and E. Halperin, 2012 A model-based approach for analysis of spatial structure in genetic data. Nat Genet 44: 725–731.

Zhang, F., M. Flickinger, S. A. G. Taliun, P. P. G. C. In, G. R. Abecasis et al., 2020 Ancestry-agnostic estimation of DNA sample contamination from sequence reads. Genome Res 30: 185–194.

